# Spatial MALDI-MSI Reveals a Coordinated Vicious Cycle of Oxidative Membrane Damage and Ceramide-Driven Sphingolipid Dysregulation in the Chronically Neuroinflamed Brain

**DOI:** 10.64898/2026.07.14.738110

**Authors:** Kent Brown, Brayden Storey, Jacob Williams, Derrick Simet, Muhammad Bilal Umar, Emma Madsen, Zhiyang Shan, Lanrong Bi

**Author notes:** Corresponding Author, Lanrong Bi - Department of Chemistry, Michigan Technological University, Houghton, Michigan 49931, Health Research Institute, Michigan Technological University, Houghton, Michigan 49931. Author Contributions. These authors contributed equally (Kent Brown, Brayden Storey, Jacob Williams).

## Abstract

**Background:** Chronic neuroinflammation is a major driver of cognitive decline, vascular cognitive impairment, and Alzheimer’s disease. However, the spatial lipidomic alterations underlying neuroinflammatory brain injury remain poorly defined. Oxidative stress and sphingolipid dysregulation have been implicated, but their regional distribution and interplay in the brain are not well characterized.

**Methods:** We performed positive-ion mode matrix-assisted laser desorption/ionization mass spectrometry imaging (MALDI-MSI) on coronal brain sections from middle-aged spontaneously hypertensive rats (SHR), a model of chronic neuroinflammation, and normotensive Wistar-Kyoto (WKY) controls. Spatial distributions and relative abundances of multiple lipid classes, including phosphatidylcholines (PCs), sphingomyelins (SMs), hexosylceramides (HexCers), ceramides, phosphatidylserines (PSs), phosphatidylinositols (PIs), phosphatidylethanolamines (PEs), phosphatidic acids (PAs), and sulfatides, were mapped and compared between genotypes. Region-of-interest analysis was used to quantify changes across cortex, hippocampus, and white-matter tracts.

**Results:** SHR brains exhibited a coordinated lipidomic signature characterized by pronounced oxidative stress and membrane remodeling. Oxidized and short-chain PCs were markedly upregulated (up to 11.6-fold), while major structural diacyl PCs were broadly downregulated. Concurrently, sphingolipids were significantly altered, with robust upregulation of SM(d36:1) (7.5-fold) and multiple HexCer species (1.5-1.9-fold), accompanied by accumulation of ceramides. These changes were accompanied by heterogeneous redistribution of PS, PI, and PE species, particularly within the hippocampus. Sulfatide patterns in white-matter tracts were also altered, suggesting myelin remodeling. Region-of-interest analysis confirmed that the most pronounced lipid alterations were concentrated in the hippocampus and white-matter regions.

**Conclusions:** Chronic neuroinflammation induces a spatially organized, multi-class lipid remodeling response in the brain, driven by advanced oxidative membrane damage and a shift toward a pro-apoptotic sphingolipid profile. The convergence of these pathways creates a vicious cycle of membrane injury, mitochondrial dysfunction, and sustained neuroinflammation that is especially prominent in the hippocampus and white matter. These spatially resolved findings provide direct evidence that oxidative stress and sphingolipid dysregulation are central, interrelated mechanisms contributing to neurovascular injury and increased risk of cognitive impairment. The study highlights the power of MALDI-MSI to uncover region-specific lipid pathology and identifies potential lipid-based targets for therapeutic intervention in neuroinflammatory brain disease.

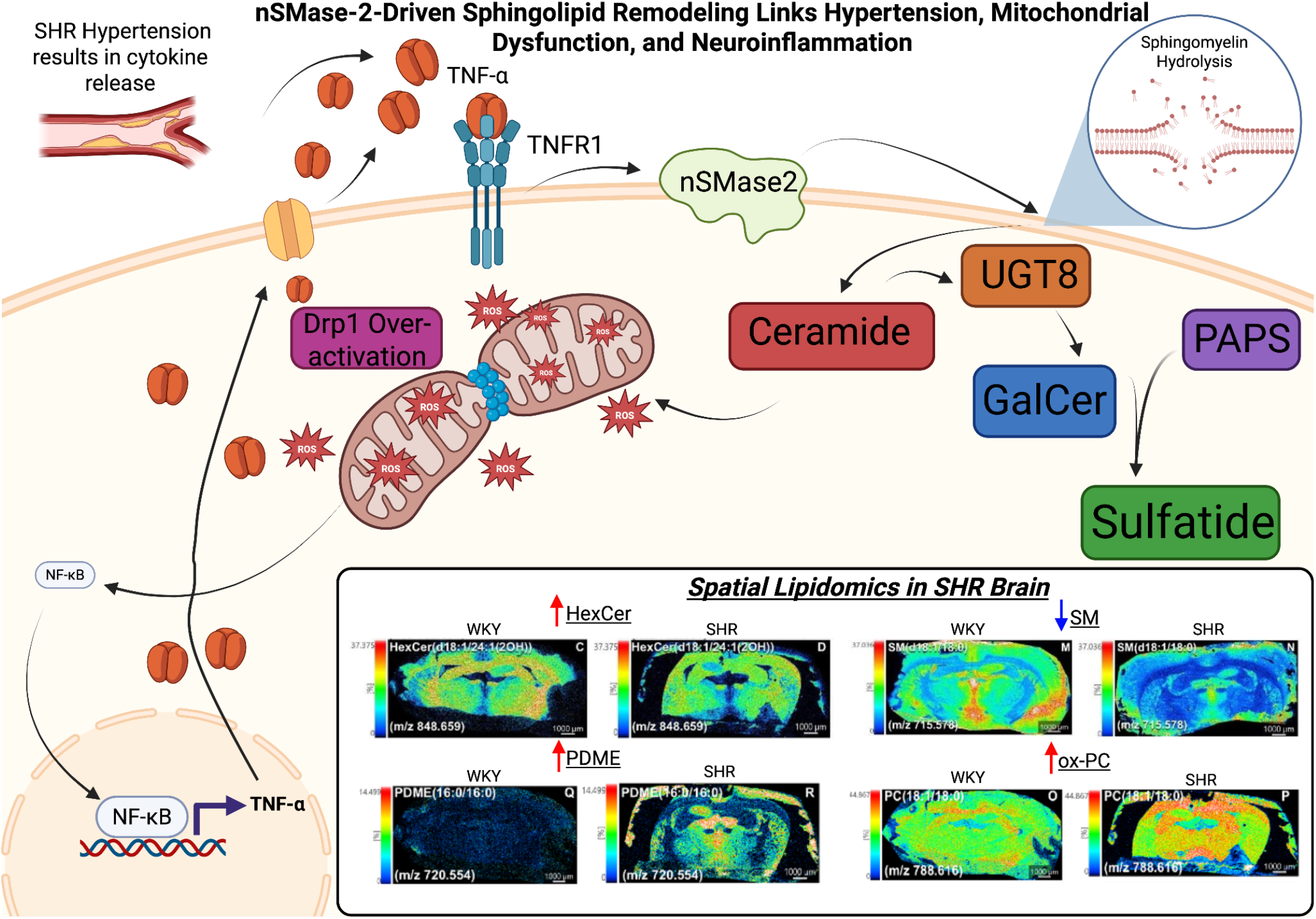

## 1. Introduction

Chronic neuroinflammation is a major driver of cognitive decline, vascular cognitive impairment (VCI), and Alzheimer’s disease (AD). Large-scale epidemiological and clinical studies have consistently demonstrated that conditions promoting sustained neuroinflammation are associated with increased risk of late-life dementia, independent of other vascular risk factors. The mechanisms underlying neuroinflammatory brain injury are multifactorial and include chronic hypoperfusion, blood-brain barrier (BBB) dysfunction, oxidative stress, and white-matter damage. Among these, oxidative stress and membrane lipid remodeling have emerged as central pathological processes that may link sustained neuroinflammatory stress to progressive neuronal and glial dysfunction [1,2].

The brain is exceptionally rich in lipids, which constitute approximately 50–60% of its dry weight[3]. Phospholipids such as phosphatidylcholines (PCs), phosphatidylethanolamines (PEs), phosphatidylserines (PSs), phosphatidylinositols (PIs), and phosphatidic acids (PAs) form the structural backbone of cellular and mitochondrial membranes, while sphingolipids including sphingomyelins (SMs), ceramides (Cers), hexosylceramides (HexCers), and sulfatides, play critical roles in membrane organization, myelin stability, and intracellular signaling. Disruption of these lipid classes can profoundly affect membrane fluidity, curvature, vesicle trafficking, and the function of membrane-embedded proteins[4,5,6]. In addition, certain lipid species, particularly ceramides and oxidized phospholipids, act as bioactive signaling molecules that can directly modulate apoptosis, inflammation, and mitochondrial function[5,6].

Oxidative stress is a well-documented feature of chronic neuroinflammation[7]. Elevated reactive oxygen species (ROS) production in the brain can initiate lipid peroxidation, leading to the formation of truncated oxidized phospholipids and reactive aldehydes such as 4-hydroxynonenal[8,9]. These electrophilic species can covalently modify proteins and nucleic acids, thereby extending oxidative damage beyond lipids and contributing to cellular dysfunction[9]. At the same time, oxidative stress can activate sphingomyelinase, resulting in the hydrolysis of sphingomyelin and accumulation of ceramide[10]. The resulting shift in the ceramide/sphingosine-1-phosphate (S1P) rheostat promotes apoptosis, mitochondrial dysfunction, and pro-inflammatory signaling while impairing protective endothelial functions[11]. This dual action of oxidative stress on both structural phospholipids and sphingolipid metabolism creates a potential vicious cycle of membrane injury and cellular stress that may be particularly detrimental in brain regions with high metabolic demand or limited antioxidant capacity[12].

Despite growing recognition of lipid alterations in neuroinflammatory brain injury, most previous studies have relied on bulk lipid extraction methods, which average lipid content across heterogeneous brain regions and cell types[12,13,14]. This approach obscures important spatial information and limits the ability to identify region-specific vulnerabilities. The hippocampus and white-matter tracts, for example, are known to be particularly susceptible to neuroinflammatory damage, yet conventional methods cannot readily distinguish whether lipid changes are uniformly distributed or concentrated in these vulnerable areas[15,16]. Furthermore, the interplay between oxidative phospholipid damage and sphingolipid dysregulation has not been systematically examined at the spatial level in models of chronic neuroinflammation.

Matrix-assisted laser desorption/ionization mass spectrometry imaging (MALDI-MSI) has emerged as a powerful technology capable of mapping the spatial distribution of hundreds of lipids directly from tissue sections while preserving anatomical context[17]. By combining high spatial resolution with molecular specificity, MALDI-MSI enables the identification of region-specific lipid alterations that may be critical for understanding the pathophysiology of complex brain diseases. This technology is particularly well suited for studying neuroinflammatory brain injury, where regional differences in oxidative burden, metabolic demand, and cellular composition are likely to produce spatially organized lipid changes[18].

The spontaneously hypertensive rat (SHR) is a well-established genetic model that recapitulates many features of chronic neuroinflammation and associated brain injury, including oxidative stress, BBB dysfunction, white-matter damage, and cognitive impairment. When compared with its normotensive counterpart, the Wistar-Kyoto (WKY) rat, the SHR provides a robust platform for investigating neuroinflammation-associated lipid alterations in the brain[19,20,21].

In our present study, we employed positive-ion mode MALDI-MSI to perform a comprehensive spatial lipidomic analysis of coronal brain sections from middle-aged SHR and WKY rats. We systematically examined alterations across multiple lipid classes, including sphingomyelins, hexosylceramides, phosphatidylcholines, phosphatidylethanolamines, phosphatidylinositols, phosphatidylserines, phosphatidic acids, sulfatides, and ceramides, as well as selected small metabolites. Our primary objectives were to (1) identify the dominant lipidomic signatures associated with neuroinflammatory brain injury, (2) characterize the spatial distribution of these changes across cortex, hippocampus, and white-matter tracts, and (3) integrate oxidative stress-related phospholipid alterations with sphingolipid dysregulation to develop a mechanistic framework linking chronic neuroinflammation to neurovascular and cognitive impairment.

We hypothesize that chronic neuroinflammation would induce a coordinated, region-specific remodeling of brain lipids, characterized by advanced oxidative damage to structural phospholipids and a shift toward a pro-apoptotic sphingolipid profile. We further propose that these lipid changes would be particularly pronounced in the hippocampus and white matter, since these regions are known to be vulnerable to neuroinflammatory injury and strongly implicated in cognitive decline. By leveraging the spatial resolving power of MALDI-MSI, our present study aimed to move beyond descriptive lipid profiling and generate mechanistic insights that could inform future therapeutic strategies targeting lipid metabolism in neuroinflammation-related brain disease. The following results detail the lipidomic alterations observed across major phospholipid and sphingolipid classes, highlight region-specific patterns of injury, and provide an integrated interpretation of how oxidative stress and sphingolipid dysregulation may converge to drive neuroinflammatory brain pathology.

## 2 Results

### 2.1.1. Overall Lipidomic Signature in the SHR Hypertensive Brain

Figure 1 provides a global overview of major lipid class alterations detected by positive-ion mode MALDI-MSI in coronal brain sections from WKY normotensive controls (left) versus SHR hypertensive rats (right). The most striking and coordinated lipidomic response in the SHR brain is pronounced oxidative stress coupled with membrane damage and sphingolipid dysregulation favoring a pro-apoptotic and pro-inflammatory state.

**Figure 1.**
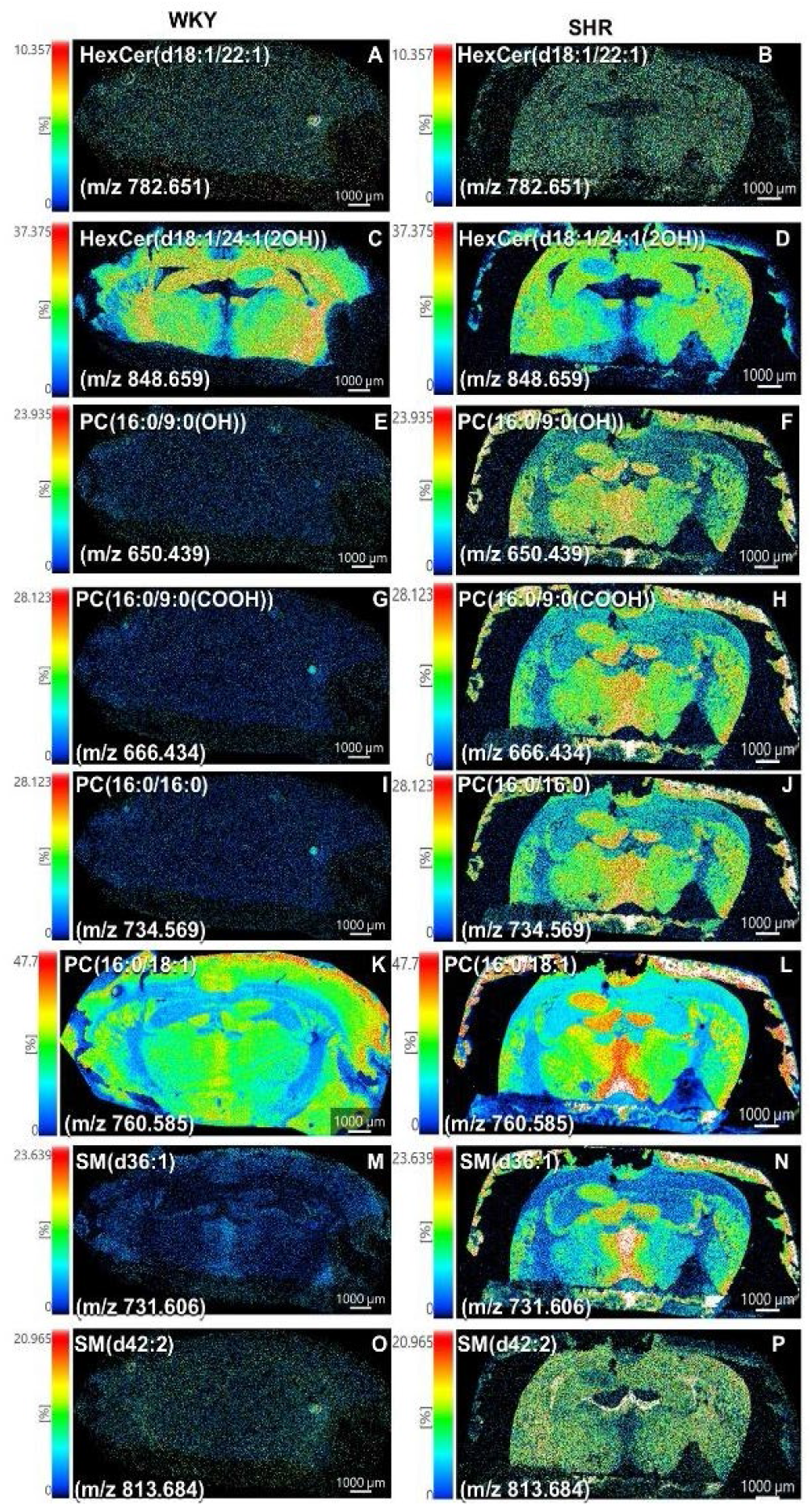
MALDI mass spectrometry imaging overview of major lipid class alterations in coronal brain sections from WKY normotensive control (left column) and SHR hypertensive (right column) rats. Representative ion images show hexosylceramides (HexCer), oxidized/short-chain phosphatidylcholines (ox-PCs), major structural phosphatidylcholines (PCs), and sphingomyelins (SM). Oxidized PCs were dramatically upregulated in SHR brains (PC(16:0/9:0(COOH)) 11.58-fold, PC(16:0/9:0(OH)) 8.07-fold, PC(18:0/9:0(OH)) 8.45-fold), while major structural PCs were strongly downregulated. Sphingolipids showed robust increases, including SM(d36:1) (7.51-fold) and HexCer species (1.5-1.9-fold). Color scale = relative ion intensity (% of maximum, normalized across panels). Scale bar = 1000 µm. Data acquired in positive-ion mode using DAN matrix. These images highlight the two dominant lipidomic signatures in the hypertensive brain: pronounced oxidative stress (oxidized PCs) and sphingolipid dysregulation (SM and HexCer upregulation).

### 2.1.2. Evidence of Advanced Oxidative Stress and Lipid Peroxidation

Oxidized and short-chain phosphatidylcholines were markedly upregulated in SHR brains. The largest increases were observed for: PC(16:0/9:0(COOH))(11.58-fold), PC(16:0/9:0(OH))(8.07-fold), PC(18:0/9:0(OH)) (8.45-fold). These truncated oxidized species are direct products of reactive oxygen species (particularly hydroxyl radicals) attacking polyunsaturated fatty acyl chains, indicating that lipid peroxidation has progressed beyond hydroperoxide formation to chain cleavage. This generates reactive aldehydes (e.g., 4-hydroxynonenal) capable of covalently modifying proteins and DNA, thereby amplifying cellular injury.

### 2.1.3. Membrane Phospholipid Depletion and Remodeling

Concomitantly, major structural diacyl PCs (including PC(16:0/16:0), PC(16:0/18:1), and their alkali adducts) were substantially downregulated (0.18–0.91-fold). This depletion indicates that abundant membrane phospholipids are being consumed during oxidative damage or are undergoing accelerated remodeling/turnover as cells attempt to preserve membrane integrity. Together, the reciprocal changes in oxidized vs. structural PCs reflect widespread oxidative injury to cellular and mitochondrial membranes, a central pathological feature of hypertensive brain injury.

### 2.1.4. Sphingolipid Dysregulation and Shift Toward a Pro-Apoptotic/Pro-Inflammatory State

Sphingolipids were consistently upregulated in SHR tissue, most notably: SM(d36:1)(7.51-fold), Multiple HexCer species (1.5–1.9-fold). When considered together with the elevation of ceramides (shown in later figures), these changes point to heightened sphingomyelinase activity (stimulated by ROS and oxidized lipids) and/or increased *de novo* synthesis, resulting in ceramide accumulation. This shifts the ceramide/S1P rheostat toward a pro-apoptotic, pro-inflammatory, and vasoconstrictive state while impairing protective endothelial functions. The concurrent rise in HexCer (and alterations in sulfatides) further suggests dysregulation of glycosphingolipid metabolism that may compromise myelin stability and oligodendrocyte function.

### 2.1.5. Spatial Distribution and Regional Vulnerability

These lipid alterations were spatially distributed across cortex, hippocampus, and white-matter tracts, indicating that hypertensive brain injury affects both gray and white matter and involves neuronal as well as glial compartments. The hippocampus and white matter, regions particularly vulnerable to hypertension, are strongly linked to cognitive decline and vascular cognitive impairment (VCI). The spatial consistency of the changes supports the conclusion that lipid remodeling represents a fundamental, brain-wide response to chronic hypertensive stress rather than a localized phenomenon.

### 2.2.1. Spatial lipidomic alterations in the brains of spontaneously hypertensive rats (SHR) versus normotensive WKY controls revealed by MALDI-MSI

Positive-ion mode MALDI mass spectrometry imaging (MALDI-MSI) was performed on coronal brain sections from WKY normotensive and SHR hypertensive rats using 1,5-diaminonaphthalene (DAN) matrix. A curated set of 29 lipid species across major classes (primarily phosphatidylcholines (PCs), sphingomyelins (SMs), hexosylceramides (HexCers), and selected phosphatidylethanolamines (PEs)) was extracted from the imaging data matrix for comparative analysis (Supplementary Table 1). Relative ion intensities were compared between groups, and representative ion images are presented in Figure 2.

**Figure 2.**
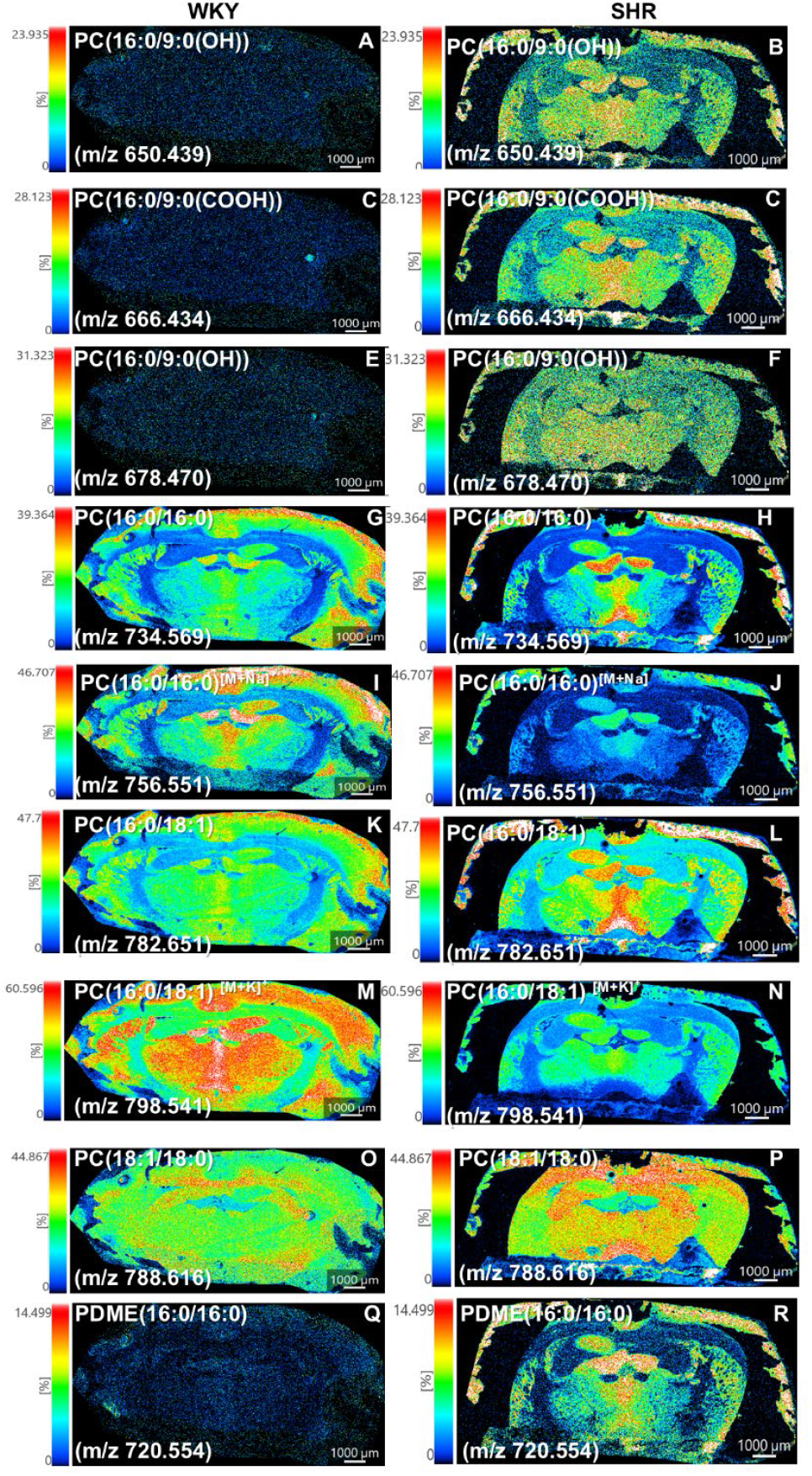
Oxidative stress and altered phosphatidylcholine metabolism in the chronically neuroinflamed rat brain. Positiveion mode MALDI-MSI of coronal brain sections from WKY (left) and SHR (right) rats. (A–C) Oxidized/short-chain PCs are strongly upregulated in SHR, indicating advanced lipid peroxidation. (D–H) Major structural diacyl PCs are markedly downregulated, consistent with membrane remodeling. (Q-R) PDME(16:0/16:0), a key precursor in PC biosynthesis, is robustly upregulated (∼7.35-fold), suggesting disrupted phospholipid methylation. These reciprocal changes demonstrate coordinated dysregulation of phosphatidylcholine metabolism under chronic neuroinflammation. Scale bar = 1000 µm.

### 2.2.2. Oxidized and truncated PCs are strongly upregulated in SHR brains

The most prominent changes occurred within the PC class. Oxidized/short-chain PC species were markedly elevated in SHR compared with WKY controls. PC(16:0/9:0(COOH)) (m/z 666.4341) showed an ∼11.6-fold increase, PC(16:0/9:0(OH)) (m/z 650.4392) an ∼8.1-fold increase, and PC(18:0/9:0(OH)) (m/z 678.4705) an ∼8.4-fold increase in SHR (Figure 2A–C). MSI images revealed that these oxidized PCs exhibited intense, regionally enriched signals (warm colors) predominantly in cortical layers and hippocampal formations of SHR sections. In contrast, WKY sections displayed diffuse, low-intensity signals (cool colors) with more uniform distribution across the parenchyma.

### 2.2.3. Major structural diacyl PCs are downregulated in SHR

In contrast, multiple abundant diacyl PC species were significantly reduced in SHR brains. PC(16:0/16:0) adducts (m/z 734.5694 [M+H]^+^, 756.5514 [M+Na]^+^, 772.5253 [M+K]^+^, 713.4518 [M+TMA]^+^) exhibited fold changes ranging from 0.18 to 0.65. PC(16:0/18:1) species (m/z 760.5851, 782.567, 798.541) showed fold changes of 0.29–0.83. Additional downregulated PCs included PC(38:4), PC(38:6), PC(40:6), and PC(18:1/18:0) variants (fold changes 0.31–0.61).

Corresponding MSI images (Figure 3D–F) demonstrated globally reduced signal intensity in SHR sections, with particularly noticeable attenuation in white matter tracts and subcortical regions. WKY images retained robust, widespread distribution of these structural PCs throughout gray and white matter.

**Figure 3.**
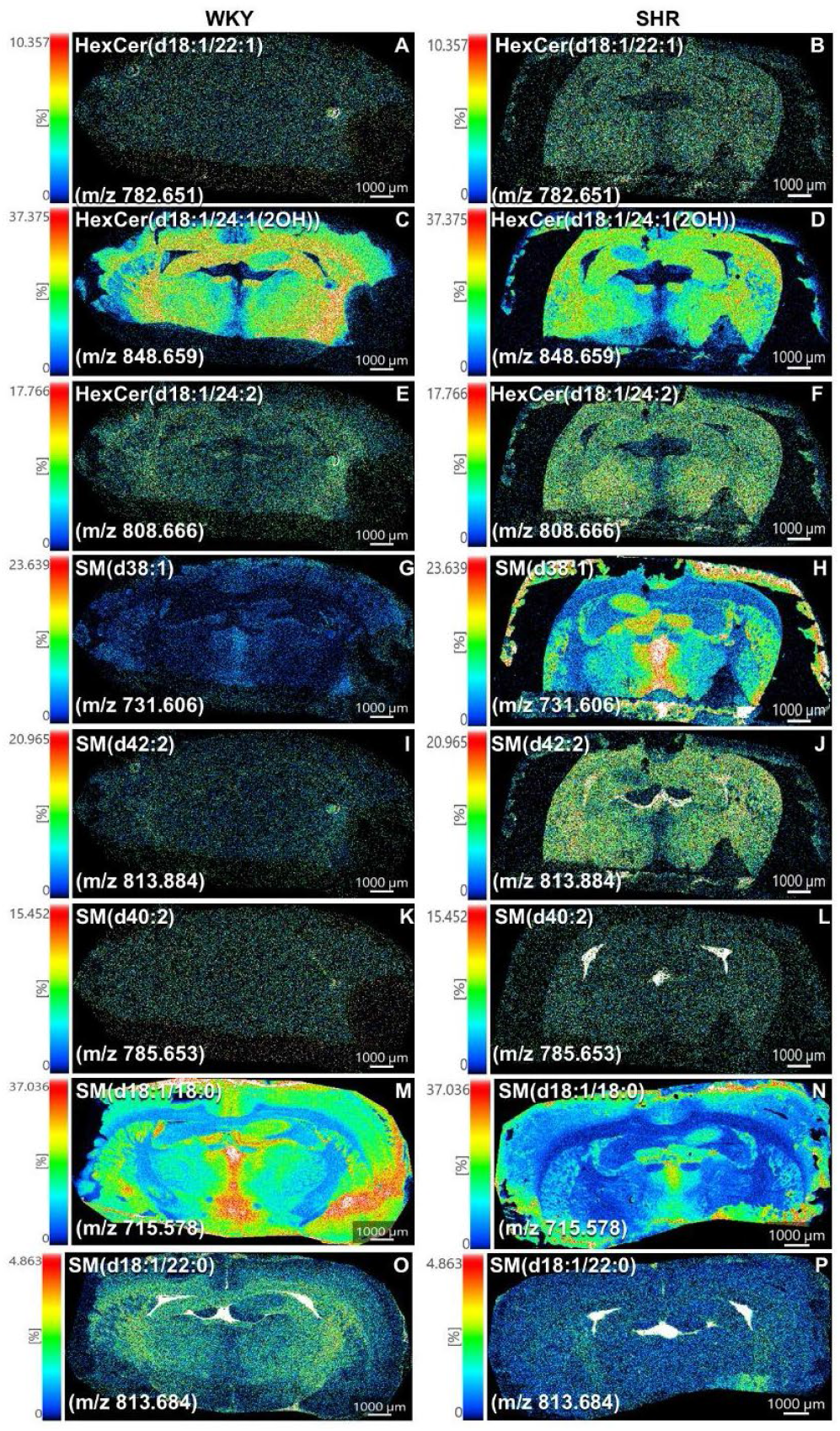
Positive-ion mode MALDI-MSI reveals sphingolipid dysregulation favoring a pro-apoptotic state in the chronically neuroinflamed rat brain. Coronal brain sections from normotensive WKY control rats (left) and SHR rats (right) were analyzed by MALDI-MSI in positive-ion mode with DAN matrix. Panels show side-by-side ion images (WKY left, SHR right) with independent color scales (0–100% relative intensity, normalized). Scale bar = 1000 μm. (A–D) Hexosylceramides (HexCer) are upregulated in SHR brains, indicating altered glycosphingolipid metabolism. Representative species include HexCer(d18:1/22:1) (*m/z* 782.650), HexCer(d18:1/24:2) (*m/z* 808.666), and HexCer(d18:1/24:1(2OH)) (*m/z* 848.659), with multiple long-chain variants showing 1.5–1.9-fold increases. (E–H) Sphingomyelins (SM) exhibit robust upregulation, consistent with enhanced sphingomyelinase activity. Key species include SM(d36:1) at *m/z* 731.606 (7.5-fold increase) and SM(d42:2) at *m/z* 813.684 (4.5-fold increase).

### 2.2.4. Sphingolipids and selected PE species show differential regulation

Sphingomyelin SM(d36:1) (m/z 731.6062) was upregulated ∼7.5-fold in SHR, with MSI images revealing enhanced, somewhat punctate signals in discrete anatomical structures (Figure 2G). HexCer(d18:1/24:2) (m/z 808.6661) showed ∼1.9-fold elevation. SM(d42:2) and certain HexCer species (e.g., HexCer(d18:1/22:1)) displayed moderate increases (1.1–4.5-fold range).

Among PEs, PE(18:0/22:6) (m/z 792.5538) was elevated ∼2.65-fold in SHR, while other PE species showed more modest or variable changes. MSI images for SM and HexCer classes (Figure 2G–H) highlighted region-specific intensity differences between genotypes, often more pronounced in cortical and periventricular areas in SHR.

### 2.2.5. Quantitative ROI analysis confirms visual MSI observations

Region-of-interest (ROI) intensity measurements extracted from the imaging data matrix (Figure 2I) quantitatively validated the visual patterns observed in the ion images. Mean intensities for oxidized PCs were substantially higher in SHR, while major diacyl PCs showed consistent reductions. These quantitative trends aligned closely with the spatial distributions captured in the MSI panels. These results demonstrate profound lipidomic remodeling in the SHR brain, characterized by elevated oxidative stress-associated PC species, reduced structural membrane PCs, and altered sphingolipid and PE profiles, with clear spatial heterogeneity between WKY and SHR genotypes.

### 2.3.1. Robust Upregulation of Sphingomyelins in the SHR Hypertensive Brain

MALDI-MSI analysis of coronal brain sections revealed a striking and consistent upregulation of sphingomyelins (SM) in SHR hypertensive rats compared with WKY normotensive controls. The most prominent change was observed for SM(d36:1) at *m/z* 731.606, which showed a 7.51-fold increase in SHR brain tissue. SM(d42:2) at *m/z* 813.684 was also markedly elevated (4.47-fold increase). Other SM species, including SM(d40:2), exhibited moderate but consistent upregulation.

Ion images demonstrated that the elevated SM signals in SHR brains were distributed across multiple brain regions, with particularly intense labeling in the cortex and white-matter tracts. In contrast, WKY control sections showed relatively low and diffuse SM signals. These findings indicate that chronic hypertension is associated with substantial accumulation of sphingomyelins, especially long-chain and very-long-chain species.

### 2.3.2. Upregulation of Hexosylceramides and Dysregulation of Glycosphingolipid Metabolism

In parallel with SM changes, several hexosylceramide (HexCer) species were significantly upregulated in SHR brains. HexCer(d18:1/22:1) at *m/z* 782.650, HexCer(d18:1/24:2) at *m/z* 808.666, and HexCer(d18:1/24:1(2OH)) at *m/z* 848.659 showed 1.5-to 1.9-fold increases relative to WKY controls.

The spatial distribution of HexCer signals in SHR sections was more intense and regionally heterogeneous than in WKY brains, with notable enrichment in white-matter regions. These alterations suggest dysregulation of glycosphingolipid metabolism, which may compromise myelin stability and oligodendrocyte function under hypertensive conditions.

### 2.3.3. Sphingolipid Remodeling as a Central Feature of Hypertensive Brain Pathology

The coordinated upregulation of sphingomyelins and hexosylceramides observed in Figure 2 represents a major lipidomic signature of the SHR hypertensive brain and provides important mechanistic insight when integrated with the oxidative stress findings from Figures 1 and 2.

Elevated SM levels, particularly the robust increase in SM(d36:1), are consistent with increased sphingomyelinase activity stimulated by reactive oxygen species and oxidized lipids. This leads to enhanced hydrolysis of sphingomyelin and subsequent ceramide accumulation (further supported by data in Figure 9). The resulting shift in the ceramide/S1P rheostat promotes apoptosis, mitochondrial dysfunction, inflammation, and vasoconstriction while impairing protective endothelial nitric oxide signaling.

Concurrently, the upregulation of HexCer species points to altered glycosphingolipid metabolism that may disrupt myelin integrity and oligodendrocyte homeostasis. Given that white-matter tracts showed particularly strong HexCer and SM signals in SHR brains, these changes likely contribute to myelin damage and white-matter injury, which are well-recognized features of hypertensive brain pathology and vascular cognitive impairment.

Spatially, the sphingolipid alterations were not confined to a single region but were distributed across the cortex, hippocampus, and white matter, indicating a brain-wide response to chronic hypertensive stress that affects both neuronal and glial compartments. The hippocampus and white matter, these regions are highly vulnerable to hypertension and exhibited pronounced changes, providing a potential lipidomic link to the cognitive decline and increased Alzheimer’s disease risk observed in hypertensive individuals. The data in Figure 3 demonstrate that hypertension induces a pro-apoptotic and pro-inflammatory sphingolipid profile that acts synergistically with oxidative membrane damage (Figures 1 and 2). This creates a vicious cycle of membrane injury, mitochondrial dysfunction, neuroinflammation, and myelin compromise. These spatially resolved findings strongly support the hypothesis that sphingolipid dysregulation is a central, interrelated mechanism driving neurovascular injury and cognitive impairment in the hypertensive brain.

### 2.4.1. Distinct Regional Distribution Patterns of Phosphatidylserines in WKY and SHR Brains

MALDI-MSI analysis revealed clear differences in the spatial distribution of phosphatidylserine (PS) species between WKY normotensive control and SHR hypertensive rat brains. In WKY sections, PS ion signals were generally uniform and homogeneous across cortical layers and the hippocampus. In contrast, SHR brains exhibited more heterogeneous and regionally accentuated PS distribution, with notable variations in signal intensity between cortical subregions and the hippocampus. These patterns were consistently observed across multiple PS species detected in positive-ion mode.

### 2.4.2. Heterogeneous and Patchy PS Signals in the SHR Hippocampus and Cortex

A particularly striking feature in SHR brains was the patchy and focal nature of PS signals, especially within the hippocampus and adjacent cortical areas. While WKY controls displayed relatively even PS distribution, SHR sections frequently showed discrete hotspots and irregular signal patterns. This heterogeneity was most pronounced in the hippocampal formation, where focal areas of elevated PS intensity were observed against a background of lower signal. Similar patchy distributions were also noted in cortical regions of SHR brains, suggesting localized membrane alterations rather than global changes.

### 2.4.3. Phosphatidylserine Remodeling as a Marker of Membrane Stress and Apoptotic Signaling

The observed PS imaging patterns provide important mechanistic insight into hypertension-associated membrane remodeling. Phosphatidylserine is normally sequestered on the inner leaflet of the plasma membrane. Its redistribution or externalization is an early and critical event in apoptosis, serving as a recognition signal for microglial phagocytosis of damaged cells. The more heterogeneous and patchy PS distribution in SHR brains, particularly the focal hotspots in the hippocampus, is consistent with increased localized membrane stress, loss of phospholipid asymmetry, and elevated apoptotic activity under chronic hypertensive conditions.

These PS alterations complement and extend the oxidative stress signature documented in Figures 1 and 2. Oxidative damage to membrane phospholipids (evidenced by elevated oxidized PCs) can disrupt membrane asymmetry and promote PS externalization. The concurrent sphingolipid dysregulation (Figure 3), including ceramide accumulation, further amplifies apoptotic signaling. Thus, the PS changes observed here likely reflect an integrated cellular response to oxidative injury and sphingolipid-mediated stress in the hypertensive brain.

### 2.4.4. Hippocampal Vulnerability and Implications for Cognitive Decline

The pronounced heterogeneity of PS signals in the SHR hippocampus is particularly significant given the well-established vulnerability of this region to hypertension. The hippocampus plays a central role in learning, memory, and cognitive function, and is highly susceptible to vascular and metabolic stress. The focal PS hotspots observed in SHR sections may indicate localized apoptotic activity or membrane damage that could contribute to neuronal loss or synaptic dysfunction. These findings provide a potential lipidomic link between hypertension, hippocampal injury, and the increased risk of cognitive decline and Alzheimer’s disease pathology observed in hypertensive populations. Spatially, PS alterations were not restricted to the hippocampus but were also distributed across cortical and white-matter regions, indicating that membrane remodeling affects multiple brain compartments. The ability of MALDI-MSI to resolve these region-specific and heterogeneous PS patterns underscores the value of spatial lipidomics in uncovering subtle but biologically meaningful changes that may be averaged out in bulk tissue analyses.

The data in Figure 4 demonstrate that hypertension induces regionally heterogeneous membrane phospholipid remodeling, with the hippocampus showing particular vulnerability. When integrated with the oxidative PC damage (Figures 1–2) and sphingolipid dysregulation (Figure 3), these findings support a model in which chronic hypertensive stress drives a coordinated membrane injury response involving lipid peroxidation, loss of asymmetry, apoptotic signaling, and neuroinflammation. This membrane-centric pathology likely contributes to neurovascular dysfunction, blood-brain barrier impairment, and long-term cognitive impairment in hypertension.

**Figure 4.**
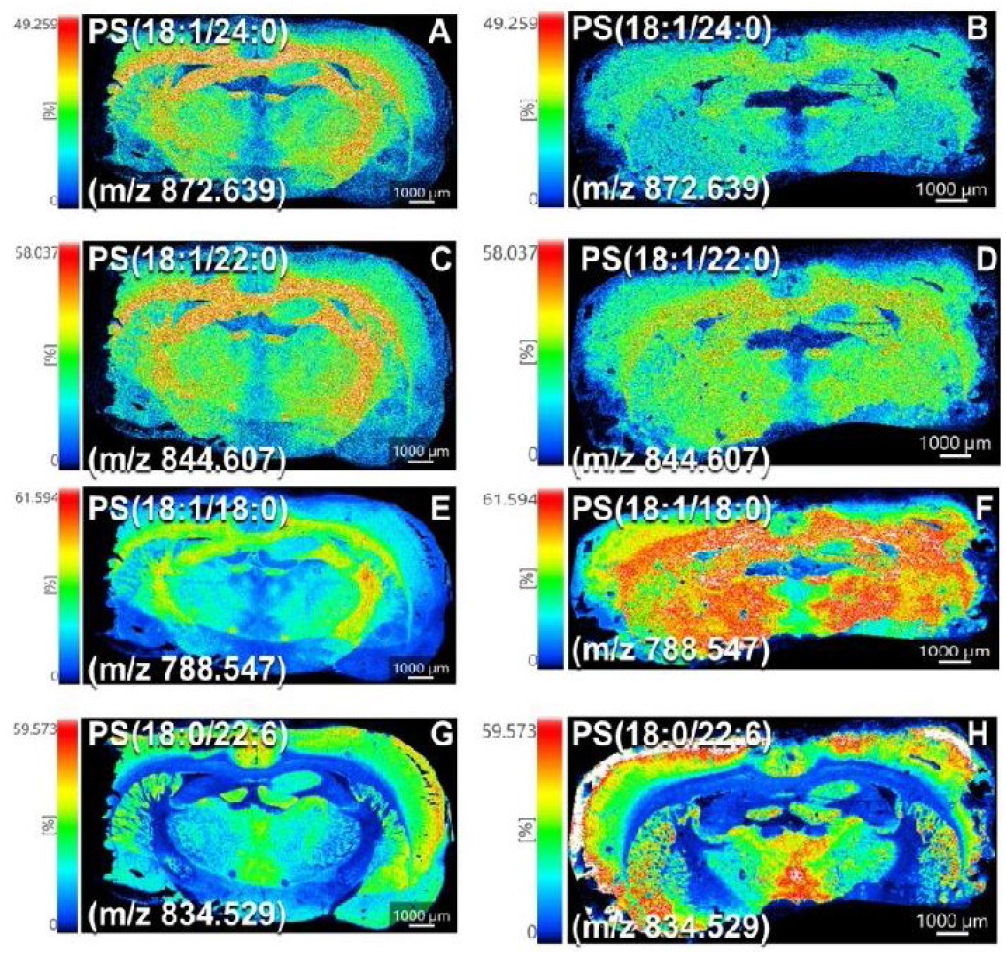
Positive-ion mode MALDI-MSI reveals region-specific and heterogeneous distribution of phosphatidylserines in the SHR hypertensive rat brain. Coronal brain sections from normotensive WKY control rats (left column) and spontaneously hypertensive rats (SHR; right column) were analyzed by MALDI-MSI in positiveion mode using DAN matrix. Panels display side-by-side ion images (WKY left, SHR right) with individual color scales (0–100% relative intensity, normalized). Scale bar = 1000 μm. Phosphatidylserine (PS) species showed distinct regional distributions with clear differences between groups. WKY controls exhibited relatively uniform PS signals across cortical and hippocampal regions, whereas SHR brains displayed heterogeneous, patchy distributions, particularly in the hippocampus and cortical layers. Several PS species were moderately upregulated or redistributed in SHR, with increased intensity, irregular patterns, focal hotspots, and layer-specific or subcortical enrichment.

### 2.5.1. Region-Specific Distribution of Phosphatidylinositols in WKY and SHR Brains

MALDI-MSI analysis of coronal brain sections revealed distinct spatial distribution patterns of phosphatidylinositol (PI) species between WKY normotensive control and SHR hypertensive rats. In WKY brains, PI ion signals were generally strong and relatively homogeneous, with clear enrichment in gray matter regions, particularly the cerebral cortex and hippocampus. In contrast, SHR brains exhibited more heterogeneous and irregular PI distribution, with noticeable variations in signal intensity across cortical layers and hippocampal subregions. These spatial differences were consistently observed across multiple PI species.

### 2.5.2. Heterogeneous Spatial Patterns and Moderate Abundance Changes in SHR Brain

A key feature in SHR sections was the more irregular and patchy distribution of PI signals compared to the relatively uniform patterns in WKY controls. Several PI species showed moderate changes in overall abundance, with some species displaying focal areas of increased or decreased intensity, particularly in the hippocampus and cortical regions. While the magnitude of abundance changes was generally smaller than those observed for oxidized PCs or sphingomyelins, the spatial reorganization of PI signals was prominent and reproducible.

### 2.5.3. Data Interpretation: Phosphatidylinositol Remodeling in the Context of Hypertensive Membrane Stress

The alterations in PI distribution and abundance observed in Figure 5 provide important insight into hypertension-associated membrane remodeling. Phosphatidylinositols are essential membrane phospholipids involved in maintaining membrane curvature, regulating vesicle trafficking, and serving as precursors for phosphoinositide signaling molecules (e.g., PI(4,5)P_2_). The more heterogeneous spatial patterns in SHR brains suggest that chronic hypertension disrupts the normal organization and metabolism of PI species.

**Figure 5.**
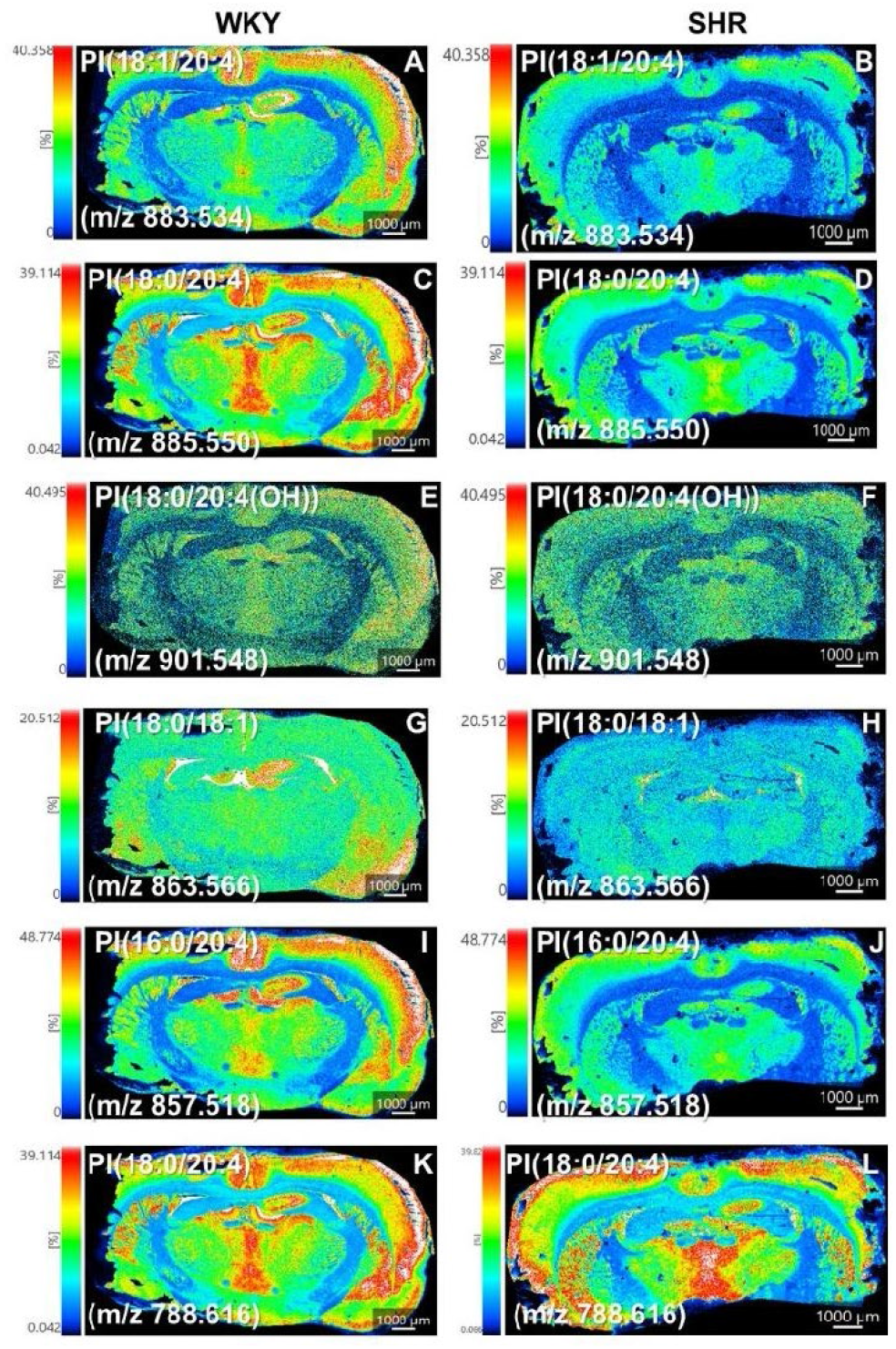
Positive-ion mode MALDI-MSI reveals region-specific alterations in phosphatidylinositol species in the chronically neuroinflamed rat brain. Coronal brain sections from normotensive WKY control rats (left) and SHR rats (right) were analyzed by MALDI-MSI in positive-ion mode with DAN matrix. Panels show side-by-side ion images (WKY left, SHR right) with independent color scales (0–100% relative intensity, normalized). Scale bar = 1000 μm. (A–D) Major phosphatidylinositol (PI) species exhibit distinct regional distributions. In WKY controls, PI signals are generally strong and relatively homogeneous, with enrichment in gray matter (cortex and hippocampus). In SHR brains, several PI species display altered intensity and increased spatial heterogeneity, including modified cortical layering and hippocampal subfield patterns. (E–H) Quantitative and spatial redistribution of PI species in SHR brains. SHR sections show moderate changes in signal intensity and more irregular, heterogeneous distributions compared with WKY, with focal variations particularly evident in the hippocampus and cortical layers. This figure demonstrates the ability of MALDI-MSI to resolve region-specific alterations in phosphatidylinositol distribution across cortex, hippocampus, and white-matter tracts in the SHR model.

These PI changes are mechanistically linked to the oxidative stress and membrane damage documented in earlier figures. Oxidative modification of membrane phospholipids (Figure 2) can alter membrane properties and influence the distribution and function of PI species. Furthermore, the sphingolipid dysregulation and ceramide accumulation observed in Figure 3 can intersect with PI signaling pathways, potentially amplifying cellular stress responses, inflammation, and apoptotic signaling. The heterogeneous PI patterns in the hippocampus are especially noteworthy, as this region is highly vulnerable to hypertensive injury and plays a central role in cognitive function.

### 2.5.4. Integration with Broader Lipidomic Changes and Pathophysiological Implications

When viewed together with the oxidative PC damage (Figures 1 and 2), sphingolipid upregulation (Figure 3), and phosphatidylserine heterogeneity (Figure 4), the PI alterations support a model of coordinated, multi-class membrane phospholipid remodeling in the hypertensive brain. This remodeling appears to involve both direct oxidative damage to structural lipids and reorganization of signaling phospholipids such as PI. The spatially resolved nature of these changes, particularly the hippocampal heterogeneity, provides a lipidomic basis for understanding how hypertension may lead to localized membrane stress, impaired signaling, and increased vulnerability to neuronal dysfunction. The findings from Figure 5 indicate that hypertension induces not only quantitative changes in phospholipid abundance but also qualitative alterations in their spatial organization. These membrane-level changes are likely to contribute to neurovascular dysfunction, blood-brain barrier impairment, and long-term cognitive decline. The ability of MALDI-MSI to detect and map these region-specific PI alterations highlights the power of spatial lipidomics to uncover subtle but biologically significant membrane changes that may be missed by conventional bulk analyses.

### 2.6.1. Region-Specific Distribution of Phosphatidylethanolamines in WKY and SHR Brains

MALDI-MSI analysis revealed clear differences in the spatial distribution of phosphatidylethanolamine (PE) species between WKY normotensive control and SHR hypertensive rat brains. In WKY sections, PE ion signals were generally strong and relatively uniform across cortical layers and the hippocampus. In contrast, SHR brains exhibited more heterogeneous and irregular PE distribution, with noticeable variations in signal intensity between cortical subregions and hippocampal areas. These spatial differences were consistently observed across multiple PE species, particularly those containing polyunsaturated fatty acyl chains.

### 2.6.2. Heterogeneous Spatial Patterns and Moderate Abundance Changes in SHR Brain

A prominent feature in SHR sections was the more patchy and irregular distribution of PE signals compared to the more homogeneous patterns in WKY controls. Several PE species, including polyunsaturated species such as PE(18:0/22:6) and PE(40:7), showed moderate changes in overall abundance accompanied by focal areas of altered intensity. These heterogeneous patterns were especially evident in the hippocampus and cortical layers of SHR brains, suggesting localized alterations in membrane phospholipid organization rather than uniform global changes.

### 2.6.3. Data Interpretation: Phosphatidylethanolamine Remodeling as Part of Coordinated Membrane Response to Oxidative Stress

The alterations in PE distribution and abundance observed in Figure 6 provide important mechanistic insight when integrated with findings from previous figures. Phosphatidylethanolamines are major structural phospholipids in cellular membranes and are particularly enriched in polyunsaturated fatty acids, making them highly susceptible to oxidative modification. The more heterogeneous spatial patterns in SHR brains are consistent with oxidative stress-induced membrane remodeling, in which PE species may undergo peroxidation, remodeling, or redistribution as cells attempt to maintain membrane integrity.

**Figure 6.**
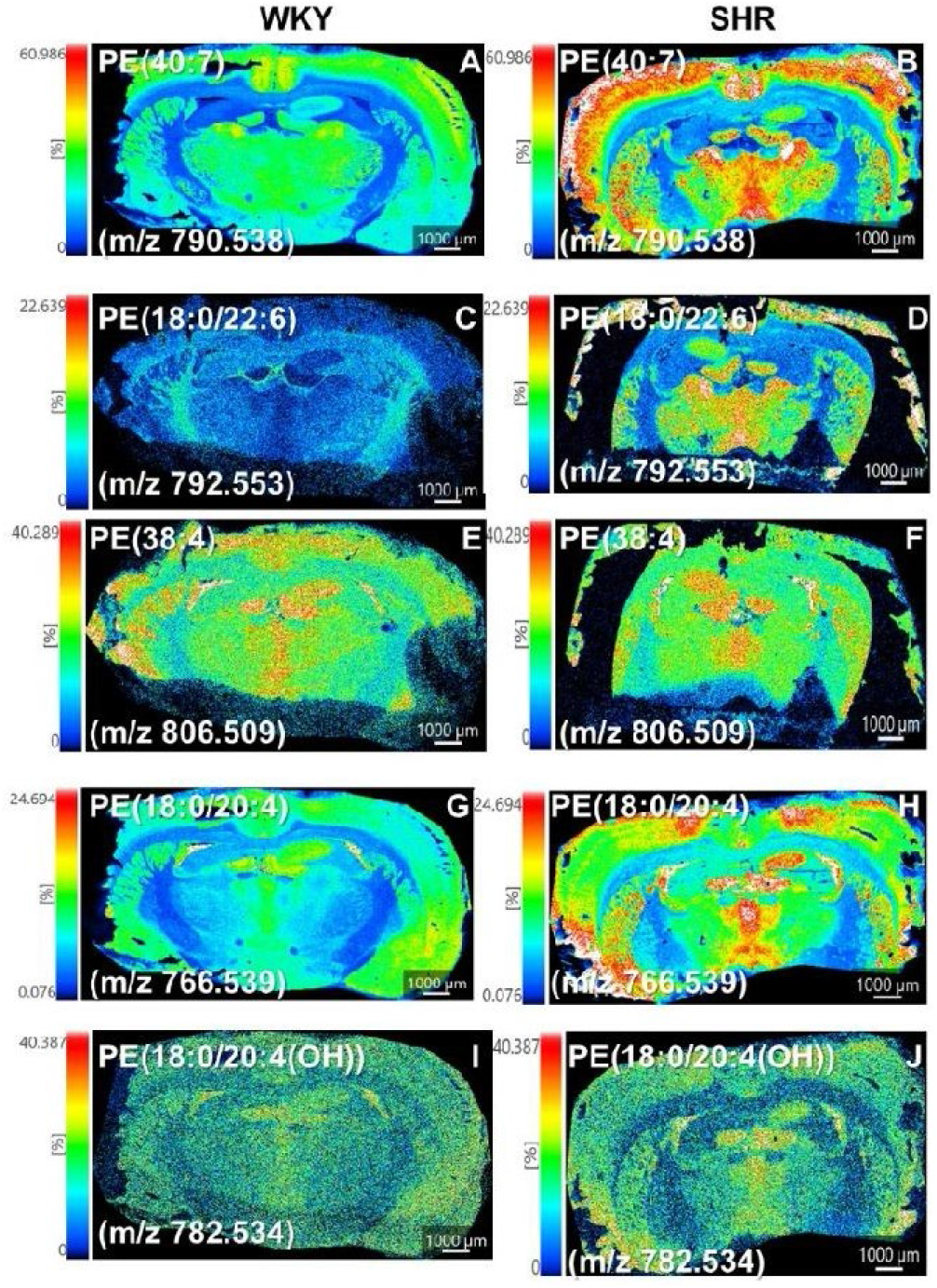
Positive-ion mode MALDI-MSI reveals region-specific alterations in phosphatidylethanolamine species in the chronically neuroinflamed rat brain. Coronal brain sections from normotensive WKY control rats (left) and SHR rats (right) were analyzed by MALDI-MSI in positive-ion mode with DAN matrix. Panels show side-by-side ion images (WKY left, SHR right) with independent color scales (0–100% relative intensity, normalized). Scale bar = 1000 μm. (A–D) Major phosphatidylethanolamine (PE) species exhibit distinct regional distributions. In WKY controls, PE signals are generally strong and relatively uniform across cortical and hippocampal regions. In SHR brains, several PE species display altered intensity and increased spatial heterogeneity, with variations in cortical layering and hippocampal subfields. (E–H) Quantitative and spatial redistribution of PE species in SHR brains. SHR sections show moderate changes in signal intensity and more irregular, heterogeneous patterns compared with WKY controls, with focal alterations particularly evident in the hippocampus and cortical layers.

These PE changes complement and extend the oxidative damage to phosphatidylcholines (Figures 1 and 2) and the alterations in phosphatidylserines and phosphatidylinositols (Figures 4 and 6). The coordinated remodeling across multiple phospholipid classes suggests that chronic hypertension imposes widespread stress on membrane structure and function. The heterogeneous PE patterns in the hippocampus are particularly significant, as this region is highly vulnerable to hypertensive injury and plays a central role in cognitive function. Focal alterations in PE may reflect localized membrane damage or adaptive remodeling responses in this vulnerable area.

### 2.6.4. Integration with Broader Lipidomic Changes and Pathophysiological Implications

When viewed together with the oxidative PC damage (Figures 1–2), sphingolipid dysregulation (Figure 3), and changes in PS and PI (Figures 4–6), the PE alterations support a model of coordinated, multi-class membrane phospholipid remodeling in the hypertensive brain. This remodeling involves both direct oxidative modification of structural lipids and reorganization of membrane composition, potentially affecting membrane fluidity, curvature, and signaling properties.

The spatially resolved nature of these PE changes, particularly in the hippocampus, provides a lipidomic basis for understanding how hypertension may lead to localized membrane stress, impaired cellular function, and increased vulnerability to neuronal injury. These findings reinforce the concept that hypertension-driven lipid remodeling is not limited to a single phospholipid class but represents a broad membrane-level response that contributes to neurovascular dysfunction, blood-brain barrier impairment, and long-term cognitive decline.

Collectively, the data in Figure 6 demonstrate that MALDI-MSI can resolve subtle, region-specific alterations in major membrane phospholipids that are difficult to detect by conventional methods. The PE changes add an important layer to the overall lipidomic signature of hypertensive brain injury and support the hypothesis that oxidative stress-induced membrane remodeling is a central mechanism linking hypertension to cognitive impairment and increased Alzheimer’s disease risk.

### 2.7.1. Region-Specific Distribution of Phosphatidic Acids in WKY and SHR Brains

MALDI-MSI analysis revealed distinct spatial distribution patterns of phosphatidic acid (PA) species between WKY normotensive control and SHR hypertensive rat brains. In WKY sections, PA ion signals were generally low to moderate and relatively uniform across cortical and hippocampal regions. In contrast, SHR brains exhibited more heterogeneous and irregular PA distribution, with noticeable variations in signal intensity between cortical layers and hippocampal subregions. These spatial differences were consistently observed across multiple PA species.

### 2.7.2. Heterogeneous Spatial Patterns and Focal Changes in SHR Brain

A prominent feature in SHR sections was the presence of focal areas of increased PA intensity against a background of lower signal, particularly in the hippocampus and cortical regions. While overall abundance changes were generally moderate compared with oxidized PCs or sphingomyelins, the spatial reorganization was clear and reproducible. SHR brains frequently displayed patchy and irregular PA patterns, in contrast to the more homogeneous distribution observed in WKY controls. These focal changes suggest localized alterations in PA metabolism or membrane dynamics under hypertensive conditions.

### 2.7.3. Phosphatidic Acid as a Marker of Membrane Remodeling and Signaling Alterations

The alterations in PA distribution observed in Figure 7 provide important mechanistic insight when integrated with findings from previous figures. Phosphatidic acids are minor but functionally significant phospholipids that serve as key intermediates in phospholipid biosynthesis and act as important signaling molecules. They are involved in regulating membrane curvature, vesicle trafficking, and cellular stress responses. The more heterogeneous and focal PA patterns in SHR brains are consistent with oxidative stress-induced membrane remodeling and potential changes in phospholipase D activity or other pathways that control PA levels.

**Figure 7.**
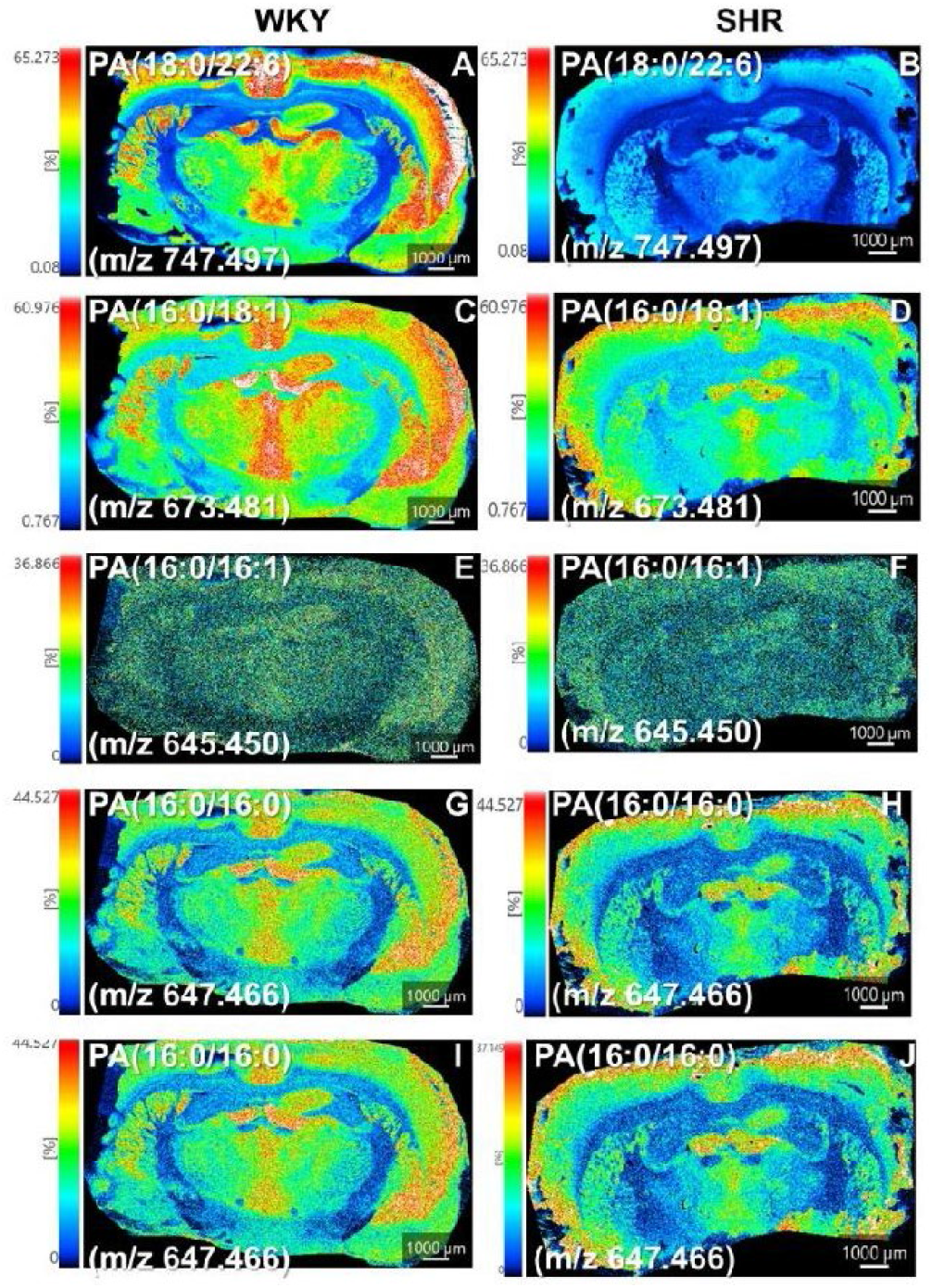
Positive-ion mode MALDI-MSI reveals region-specific alterations in phosphatidic acid species in the chronically neuroinflamed rat brain. Coronal brain sections from normotensive WKY control rats (left) and SHR rats (right) were analyzed by MALDI-MSI in positive-ion mode with DAN matrix. Panels show side-by-side ion images (WKY left, SHR right) with independent color scales (0–100% relative intensity, normalized). Scale bar = 1000 μm. (A–D) Phosphatidic acid (PA) species exhibit distinct regional distributions. In WKY controls, PA signals are generally low to moderate and relatively uniform. In SHR brains, several PA species display altered intensity and increased spatial heterogeneity, with variations across cortical layers and hippocampal subregions. (E–H) Quantitative and spatial redistribution of PA in SHR brains. SHR sections show moderate upregulation or focal redistribution, with more irregular patterns and hotspots particularly evident in the hippocampus and cortex.

These PA changes complement the oxidative damage to phosphatidylcholines (Figures 2 and 3), sphingolipid dysregulation (Figure 4), and alterations in PS, PI, and PE (Figures 4–6). The coordinated remodeling across multiple phospholipid classes, including minor signaling lipids such as PA, suggests that chronic hypertension imposes widespread stress on membrane structure and function. The focal PA hotspots observed in the hippocampus are particularly noteworthy, as this region is highly vulnerable to hypertensive injury. Such localized changes may reflect increased membrane turnover, stress signaling, or early apoptotic processes in this cognitively important area.

### 2.7.4. Integration with Broader Lipidomic Changes and Pathophysiological Implications

When viewed together with the oxidative PC damage (Figures 1–2), sphingolipid upregulation (Figure 3), and changes in PS, PI, and PE (Figures 4–6), the PA alterations support a model of coordinated, multi-class membrane phospholipid remodeling in the hypertensive brain. This remodeling involves both direct oxidative modification of structural lipids and reorganization of minor signaling phospholipids. The spatially resolved nature of these PA changes, particularly the hippocampal heterogeneity, provides a lipidomic basis for understanding how hypertension may lead to localized membrane stress and impaired cellular signaling.

Collectively, the data in Figure 8 demonstrate that hypertension affects not only major structural phospholipids but also minor signaling lipids in a region-specific manner. These findings reinforce the concept that oxidative stress-driven membrane remodeling is a central mechanism linking hypertension to neurovascular dysfunction, blood-brain barrier impairment, and long-term cognitive decline. The ability of MALDI-MSI to detect and map these subtle PA alterations highlights the power of spatial lipidomics to uncover biologically meaningful changes that may be averaged out in bulk tissue analyses.

**Figure 8.**
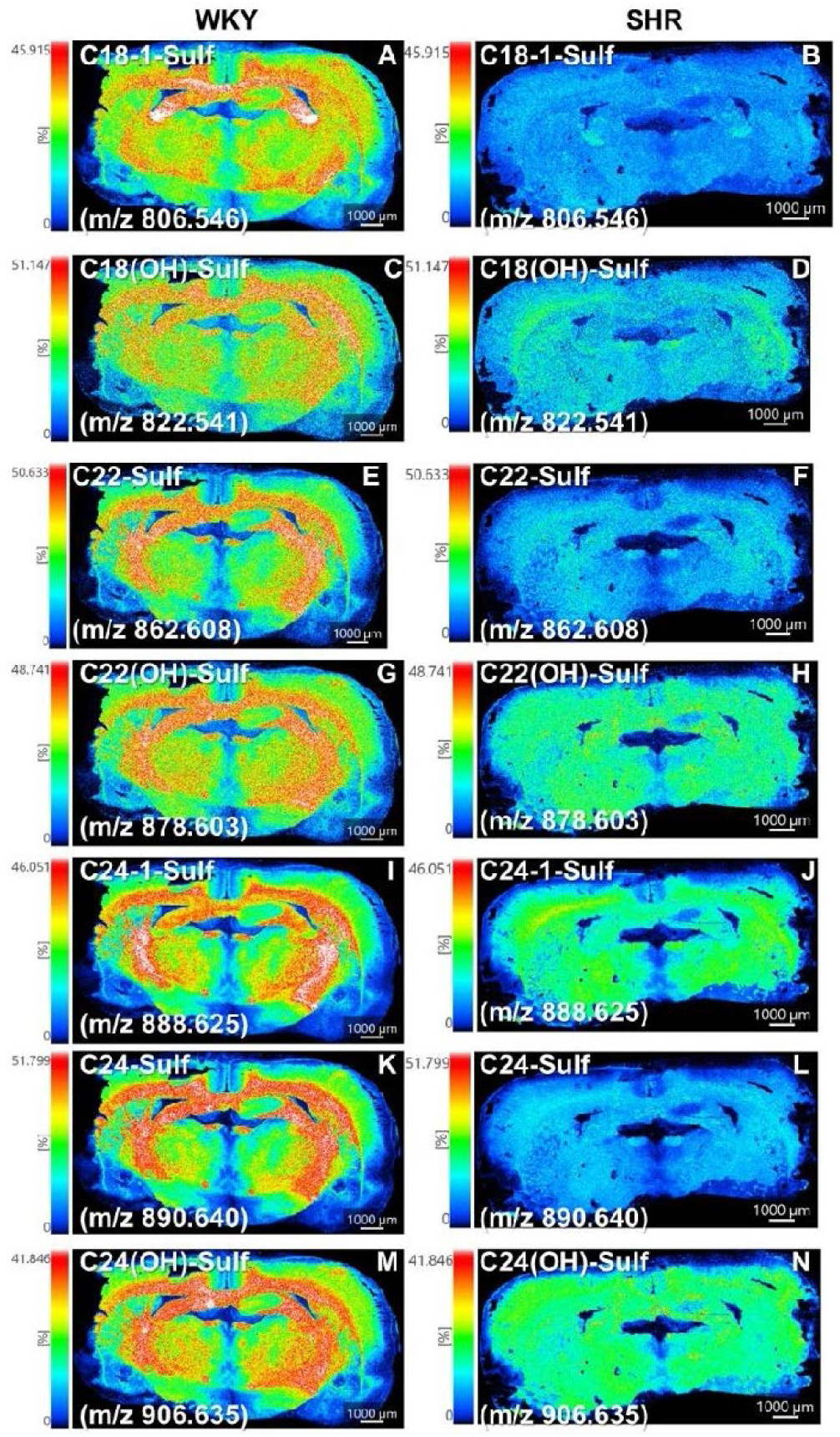
Positive-ion mode MALDI-MSI reveals altered sulfatide distribution in white-matter tracts of the chronically neuroinflamed rat brain. Coronal brain sections from normotensive WKY control rats (left) and SHR rats (right) were analyzed by MALDI-MSI in positive-ion mode with DAN matrix. Panels show side-by-side ion images (WKY left, SHR right) with independent color scales (0–100% relative intensity, normalized). Scale bar = 1000 μm. (A–D) Sulfatide species show characteristic white-matter enrichment with clear differences between groups. In both WKY and SHR brains, sulfatides are strongly enriched in white-matter tracts, consistent with their major role in myelin. SHR brains exhibit altered signal intensity and increased heterogeneity within white-matter regions compared with the more uniform distribution in WKY controls. Affected species include C18:1-, C22-, C24-, and hydroxylated sulfatides. (E–H) Quantitative and spatial redistribution of sulfatides in SHR white matter. SHR sections display moderate changes in abundance and more irregular, focal patterns within white-matter tracts.

### 2.8.1. Strong White-Matter Enrichment of Sulfatides with Differences Between WKY and SHR Brains

MALDI-MSI analysis revealed characteristic strong enrichment of sulfatide species in white-matter tracts of both WKY normotensive control and SHR hypertensive rat brains, consistent with their role as major myelin lipids. In WKY controls, sulfatide signals were generally strong and relatively uniform within white-matter regions. In contrast, SHR brains exhibited more heterogeneous and irregular sulfatide distribution within white-matter tracts, with noticeable variations in signal intensity and spatial organization compared with WKY controls. These spatial differences were consistently observed across multiple sulfatide species, including C18:1-, C22-, C24-, and hydroxylated sulfatides.

### 2.8.2. Heterogeneous Spatial Patterns and Moderate Abundance Changes in SHR White Matter

A prominent feature in SHR sections was the presence of patchy and irregular sulfatide signals within white-matter tracts, in contrast to the more homogeneous distribution observed in WKY controls. Several sulfatide species showed moderate changes in overall abundance accompanied by focal areas of altered intensity. These heterogeneous patterns were particularly evident in major white-matter bundles and subcortical regions of SHR brains, suggesting localized alterations in myelin lipid composition or stability under hypertensive conditions.

### 2.8.3. Data Interpretation: Sulfatide Alterations as Indicators of Myelin Remodeling and White-Matter Injury

The alterations in sulfatide distribution observed in Figure 8 provide important mechanistic insight when integrated with findings from previous figures. Sulfatides are essential components of myelin and play critical roles in myelin stability, oligodendrocyte function, and axon-glial interactions. The more heterogeneous and irregular sulfatide patterns in SHR white-matter tracts are consistent with oxidative stress-induced myelin remodeling or damage. This is supported by the pronounced oxidative damage to phosphatidylcholines (Figures 1 and 2) and the sphingolipid dysregulation (Figure 3), which includes ceramide accumulation that can promote oligodendrocyte stress and myelin instability. The observed sulfatide changes suggest that chronic hypertension may compromise myelin integrity through a combination of direct oxidative modification and altered sphingolipid metabolism. These findings align with the known vulnerability of white matter to hypertensive injury and provide a lipidomic basis for white-matter abnormalities frequently observed in hypertensive patients and animal models.

### 2.8.4. Integration with Broader Lipidomic Changes and Pathophysiological Implications

When viewed together with the oxidative PC damage (Figures 1–2), sphingolipid dysregulation (Figure 3), and alterations in PS, PI, PE, and PA (Figures 5–8), the sulfatide changes support a model of coordinated, multi-class membrane and myelin remodeling in the hypertensive brain. This remodeling affects both gray-matter phospholipids and white-matter myelin lipids, indicating that hypertension imposes widespread stress on neural tissue. The spatially resolved nature of these sulfatide alterations, particularly the heterogeneous patterns in white-matter tracts, provides direct evidence that myelin lipid composition is disrupted in a region-specific manner under chronic hypertensive stress. Collectively, the data in Figure 9 demonstrate that hypertension affects not only neuronal and glial membrane phospholipids but also the lipid composition of myelin. These findings reinforce the concept that oxidative stress-driven membrane and myelin remodeling is a central mechanism linking hypertension to neurovascular injury, white-matter damage, and long-term cognitive decline. The ability of MALDI-MSI to detect and map these subtle sulfatide alterations highlights the power of spatial lipidomics to uncover biologically meaningful changes in myelin that may be difficult to detect by conventional methods.

**Figure 9.**
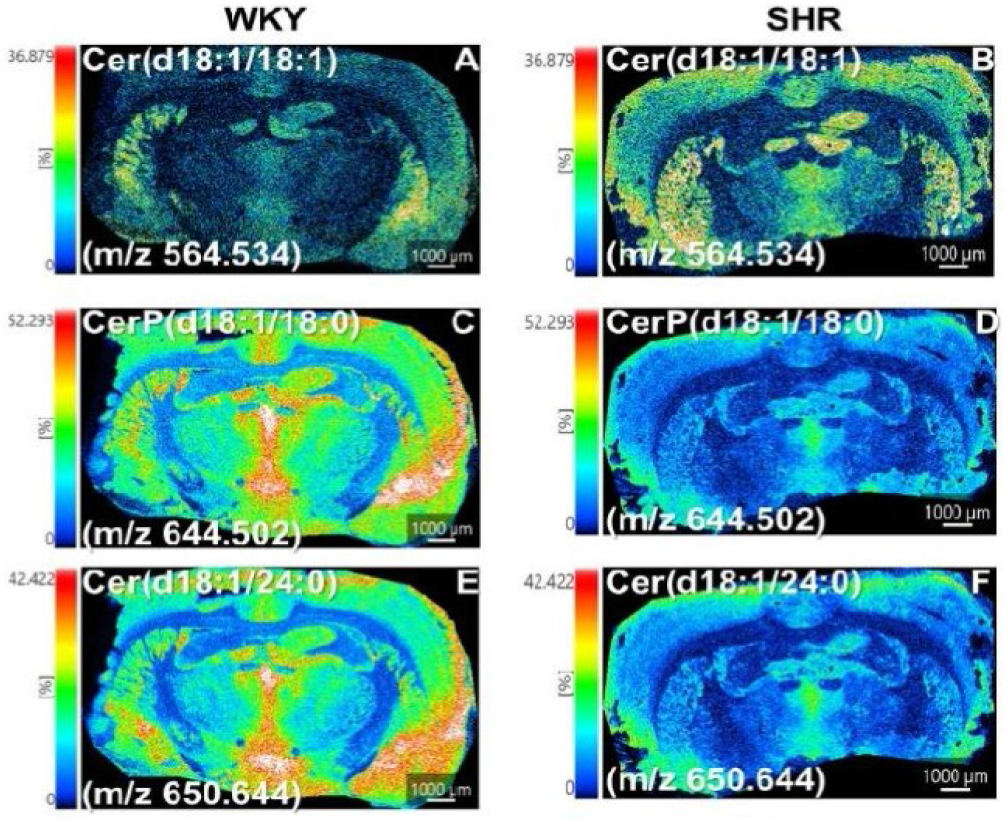
Spatial distribution of ceramide species in coronal brain sections from normotensive WKY control and hypertensive SHR rats. Positive-ion mode MALDI-MSI analysis reveals robust upregulation of multiple ceramide species in the SHR brain. Representative ion images are shown for (A, B) Cer(d18:1/18:1) (m/z 564.534), (C, D CerP(d18:1/18:0) (m/z 644.502), and (E, F) Cer(d18:1/24:0) (m/z 650.644). Left panels (A, C, E) show WKY normotensive controls, whereas right panels (B, D, F) show SHR hypertensive rats. Images are displayed using independent color scales (0-100% relative intensity, normalized within each panel). Scale bar = 1000 μm. Data acquired using 1,5-diaminoaphthalene (DAN) as the MALDI matrix. These findings demonstrate region-specific accumulation of ceramide and ceramide phosphate species in SHR brains, including prominent changes in cortical and hippocampal regions, supporting dysregulated sphingolipid metabolism and ceramide accumulation as important contributors to pro-inflammatory and stress-associated signalling in hypertensive neuroinflammation.

### 2.9.1. Upregulation of Both Short-to Medium-Chain and Long-Chain Ceramides in SHR Brain

MALDI-MSI analysis revealed robust upregulation of ceramide species in SHR hypertensive rat brains compared with WKY normotensive controls. Both short-to medium-chain ceramides (e.g., Cer(d18:1/18:0), Cer(d18:1/18:1)) and long-chain ceramides (e.g., Cer(d18:1/24:0), Cer(d18:1/24:1)) showed increased signal intensity in SHR sections. The spatial distribution in SHR brains was more heterogeneous, with notable accentuation in cortical and hippocampal regions, whereas WKY controls displayed lower and more uniform ceramide signals across these areas.

### 2.9.2. Distinct Spatial Patterns of Short/Medium-Chain versus Long-Chain Ceramides in SHR Brain

Short-to medium-chain ceramides in SHR brains showed moderate upregulation with relatively focal distribution, particularly in cortical layers and the hippocampus. Long-chain ceramides exhibited stronger and more widespread upregulation, with prominent signals in both gray-matter and white-matter regions. These differential patterns suggest that hypertension may differentially affect ceramide species with varying acyl chain lengths, potentially influencing their biophysical properties and biological activities within specific brain compartments.

### 2.9.3. Ceramide Accumulation as a Central Driver of Pro-Apoptotic and Pro-Inflammatory Signaling

The upregulation of ceramides observed in Figure 9 provides critical mechanistic insight when integrated with findings from previous figures. Ceramides are bioactive sphingolipids that act as key mediators of cellular stress responses. Their accumulation in SHR brains is consistent with increased sphingomyelinase activity (stimulated by reactive oxygen species and oxidized lipids, as shown in Figures 1 and 2) and/or enhanced *de novo* synthesis. This leads to a shift in the ceramide/S1P rheostat toward a pro-apoptotic, pro-inflammatory, and vasoconstrictive state.

Elevated ceramides promote mitochondrial dysfunction, cytochrome c release, and activation of apoptotic pathways. They also amplify inflammatory signaling and impair protective endothelial functions. The pronounced ceramide signals in the hippocampus are particularly significant, given the vulnerability of this region to hypertensive injury and its central role in cognitive function. These findings directly complement the robust upregulation of sphingomyelins and hexosylceramides (Figure 3), indicating coordinated dysregulation of the sphingolipid pathway that favors ceramide accumulation.

### 2.9.4. Integration with Broader Lipidomic Changes and Pathophysiological Implications

When viewed together with the oxidative PC damage (Figures 1-2), sphingolipid dysregulation (Figure 3), alterations in PS, PI, PE, and PA (Figures 4-7), and sulfatide changes (Figure 8), the ceramide upregulation completes the picture of widespread sphingolipid pathway remodeling in the hypertensive brain. This remodeling creates a vicious cycle in which oxidative stress drives ceramide accumulation, which in turn promotes further membrane damage, mitochondrial dysfunction, inflammation, and cell death.

Collectively, the data in Figure 9 demonstrate that hypertension induces not only quantitative increases in ceramide levels but also region-specific redistribution, particularly in cognitively vulnerable areas such as the hippocampus. These findings strongly support the hypothesis that ceramide accumulation is a central, interrelated mechanism linking oxidative stress, membrane injury, neuroinflammation, and increased risk of cognitive decline and Alzheimer’s disease pathology in hypertension. The ability of MALDI-MSI to spatially resolve these ceramide alterations provides direct evidence of their involvement in hypertensive brain injury that cannot be fully captured by bulk lipidomics.

#### 2.10. Dynamic Mathematical Modeling of the Oxidative Stress–Sphingolipid Vicious Cycle

To quantitatively integrate the spatially resolved MALDI-MSI findings and explore the dynamics of the observed lipid alterations, we developed a system of ordinary differential equations (ODEs) describing the interconnected vicious cycle between oxidative membrane damage and sphingolipid dysregulation. The model includes six state variables: oxidative stress/ROS level (**O**), oxidized/truncated phospholipids (**L**_ox), structural phospholipids (**L**_str), ceramide (**C**), sphingolipid pool (**S**), and membrane integrity/mitochondrial function (**M**). Parameters were calibrated to reproduce the key directional changes observed in the experimental dataset.

Simulations were performed for 300 arbitrary time units under low neuroinflammatory input (**I**_neuro = 0.35; WKY-like) versus elevated chronic neuroinflammatory input (**I**_neuro = 1.35; SHR-like). As shown in Figure 10, the two conditions diverged markedly over time. While the WKY simulation remained near steady-state homeostasis, the SHR simulation showed sustained increases in oxidative stress (**O**), oxidized lipids (**L**_ox), and ceramide (**C**), together with progressive depletion of structural lipids (**L**_str) and membrane integrity (**M**). The sphingolipid pool (**S**) exhibited moderate accumulation, consistent with experimental observations.

**Figure 10.**
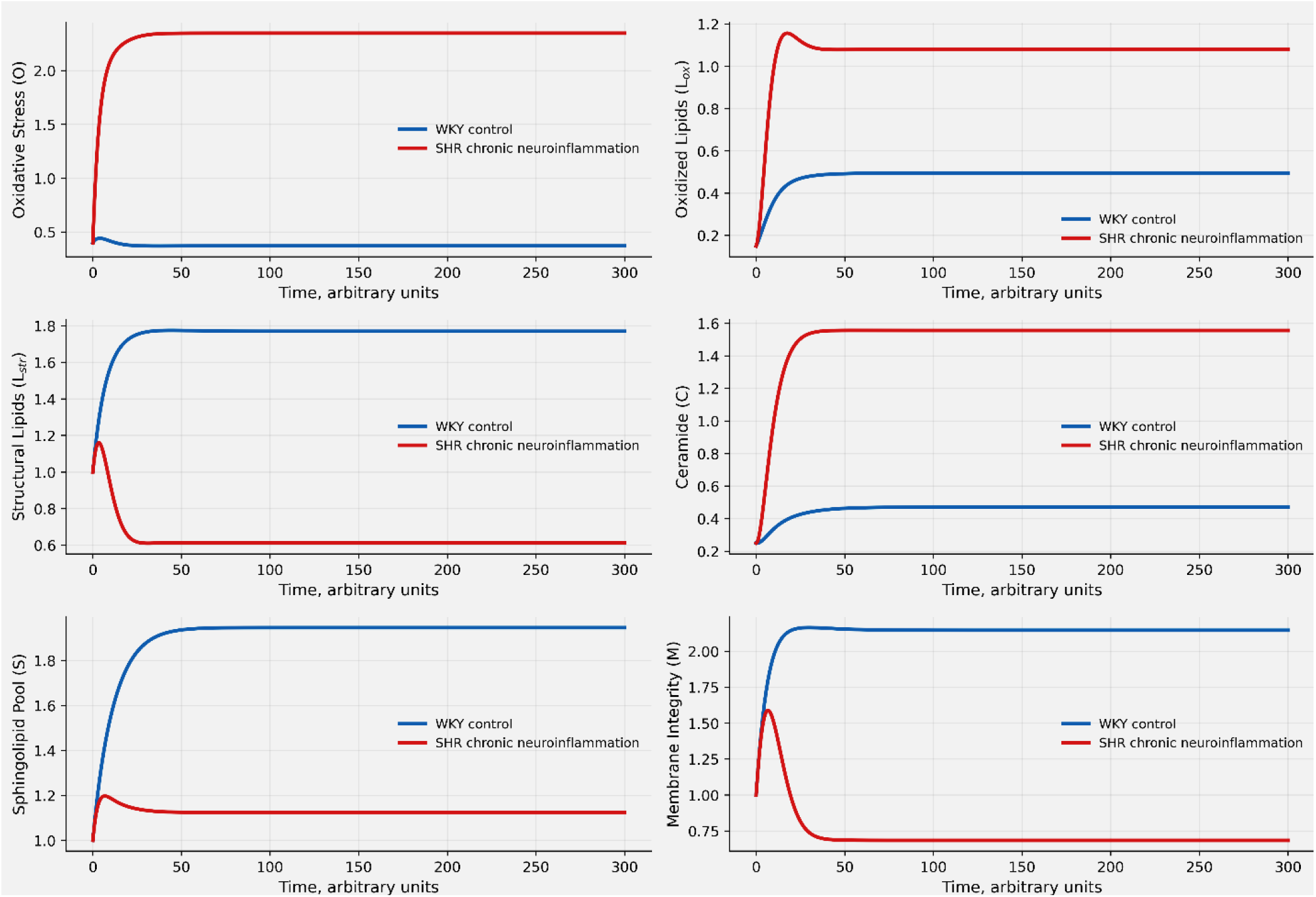
Conceptual ODE simulation of oxidative and sphingolipid remodeling in SHR versus WKY brain. Chronic neuroinflammatory drive in SHR promotes sustained oxidative stress, lipid oxidation, and ceramide accumulation, while reducing structural lipid pools and membrane integrity. The model illustrates a feed-forward pathological cycle linking oxidative lipid damage, sphingolipid remodeling, ceramide elevation, and membrane vulnerability in hypertensive neuroinflammation.

Steady-state analysis revealed fold changes that closely aligned with the MALDI-MSI data. Phase-plane analysis further indicated bistable-like behavior, suggesting that sufficiently strong and persistent neuroinflammatory drive can shift the system from a stable healthy state into a self-sustaining pathological state characterized by ongoing oxidative damage and ceramide accumulation.

The model provides mechanistic support for the experimental findings by demonstrating how elevated neuroinflammatory input can initiate a feed-forward vicious cycle: ROS-driven peroxidation of structural phospholipids generates reactive species that activate sphingomyelinase, leading to ceramide accumulation. Ceramide, in turn, promotes mitochondrial dysfunction and further membrane damage, amplifying ROS production and impairing repair mechanisms. Regional vulnerability (particularly in the hippocampus and white matter) emerges naturally when differences in metabolic demand or antioxidant capacity are considered.

Notably, the ODE framework enables in silico evaluation of therapeutic interventions. Preliminary simulations suggest that reducing ROS production or inhibiting sphingomyelinase activity can interrupt the pathological trajectory and help restore membrane integrity, with combined interventions showing greater efficacy than single-target approaches. This modeling approach thus complements the spatial lipidomic data and offers a quantitative tool for testing hypotheses and prioritizing therapeutic strategies targeting the oxidative–sphingolipid axis in chronic neuroinflammation.

## 3. Discussion

The spatial lipidomic profiling by positive-ion mode MALDI-MSI provides compelling evidence that the SHR hypertensive brain undergoes a highly orchestrated, multi-class membrane remodeling program in response to chronic neuroinflammation. This response is characterized by advanced oxidative damage to structural phospholipids, nSMase2-mediated sphingolipid reprogramming, loss of membrane asymmetry, disruption of phospholipid biosynthetic pathways, and mitochondrial bioenergetic compromise. Rather than representing isolated lipid alterations, these changes collectively establish a self-reinforcing pro-apoptotic, pro-inflammatory, and pro-oxidative milieu that preferentially affects vulnerable regions such as the hippocampus and white-matter tracts, thereby linking chronic hypertension to accelerated cognitive decline and increased risk of vascular cognitive impairment and Alzheimer’s disease/ADRD.

A central and highly reproducible finding is the pronounced accumulation of oxidized and truncated phosphatidylcholines, including PC(16:0/9:0(COOH)) (∼11.58-fold), PC(16:0/9:0(OH)) (∼8.07-fold), and PC(18:0/9:0(OH)) (∼8.45-fold), coupled with substantial depletion of major structural diacyl PCs (0.18–0.91-fold for species such as PC(16:0/16:0), PC(16:0/18:1), and their alkali adducts) (Figures 1 and 2; Supplementary Figure S2). Supplementary Figures S2 and S10 provide complete class coverage and mechanistic context, demonstrating that this reciprocal pattern reflects advanced lipid peroxidation that has progressed beyond hydroperoxide formation to chain cleavage. As detailed in Supplementary Figure S3, hydroxyl radicals abstract bis-allylic hydrogens from polyunsaturated acyl chains, initiating a cascade that generates reactive aldehydes (e.g., 4-hydroxynonenal) capable of covalently modifying proteins, DNA, and neighboring lipids [22-27]. This oxidative assault compromises membrane integrity, fluidity, and mitochondrial function, representing a primary driver of cellular injury in hypertensive neuroinflammation [28-30].

Equally critical is the robust dysregulation of the sphingolipid pathway. Significant upregulation of sphingomyelins (SM(d36:1) ∼7.51-fold, SM(d42:2) ∼4.5-fold) and multiple hexosylceramide species (1.5–1.9-fold) (Figure 3), together with accumulation of both short/medium-chain and long-chain ceramides (Figure 9), points to enhanced sphingomyelinase activity stimulated by ROS and oxidized lipids [31, 32]. Supplementary Figure S4 illustrates the resulting shift in the ceramide/S1P rheostat, while Supplementary Figure S9 details the biosynthetic framework supporting coordinated sphingolipid upregulation. Ceramide accumulation promotes Drp1-mediated mitochondrial fragmentation, cytochrome c release, NF-κB activation, and TNF-α production, thereby sustaining a vicious cycle of inflammation and mitochondrial dysfunction [33-36]. The spatial enrichment of these sphingolipids in cortex, hippocampus, and white-matter tracts further underscores their contribution to both neuronal and glial pathology, including myelin instability (Figure 8 and Supplementary Figure S8).

Spatially resolved imaging additionally reveals heterogeneous and patchy redistribution of phosphatidylserines, with focal hotspots particularly prominent in the hippocampus (Figure 4 and Supplementary Figure S1). Supplementary Figure S1 expands this analysis with long-chain and polyunsaturated PS species, confirming that the increased regional heterogeneity is a consistent feature across the PS class. Normally sequestered on the inner leaflet, PS redistribution serves as an early “eat-me” signal for microglial phagocytosis and is mechanistically linked to oxidative damage and ceramide accumulation (Supplementary Figure S5). This loss of membrane asymmetry amplifies apoptotic signaling and contributes to localized neuronal vulnerability.

Remodeling is not confined to PC and sphingolipids but extends across multiple classes. Phosphatidylinositols (Figure 5), phosphatidylethanolamines (Figure 6), and phosphatidic acids (Figure 7) exhibit moderate abundance changes accompanied by increased spatial heterogeneity, particularly in hippocampal and cortical subregions. A standout finding is the dramatic ∼7.35-fold upregulation of PDME(16:0/16:0) (Figure 2), an intermediate in the PEMT-mediated methylation pathway from PE to PC (Supplementary Figure S7). This accumulation, most pronounced in the hippocampus, suggests impaired or compensatory disruption of phospholipid biosynthesis under sustained oxidative and metabolic stress. Concurrent spatially heterogeneous alterations in energy-related metabolites (e.g., malate, glycerol-3-phosphate, creatine phosphate, arginine) further indicate mitochondrial dysfunction and impaired bioenergetics.

Figure 11 provides a comprehensive integrative model that unifies these observations around mitochondria-associated endoplasmic reticulum membrane (MAM) dysfunction. Chronic hypertensive stress disrupts MAM integrity, leading to dysregulated Ca^2+^ flux (via IP3R-GRP75-VDAC1), impaired MFN1/2-mediated tethering, Drp1 overactivation, reduced PE synthesis from PS due to phosphatidylserine decarboxylase (PSD) downregulation (Supplementary Figure S11), inner mitochondrial membrane destabilization, excessive mtROS production, cytochrome c release, and inflammasome activation. This MAM-centered cascade interconnects oxidative lipid damage, ceramide accumulation, membrane remodeling, and inflammatory signaling, creating a feed-forward loop that drives neurovascular dysfunction, white-matter injury, and progressive neurodegeneration.

**Figure 11.**
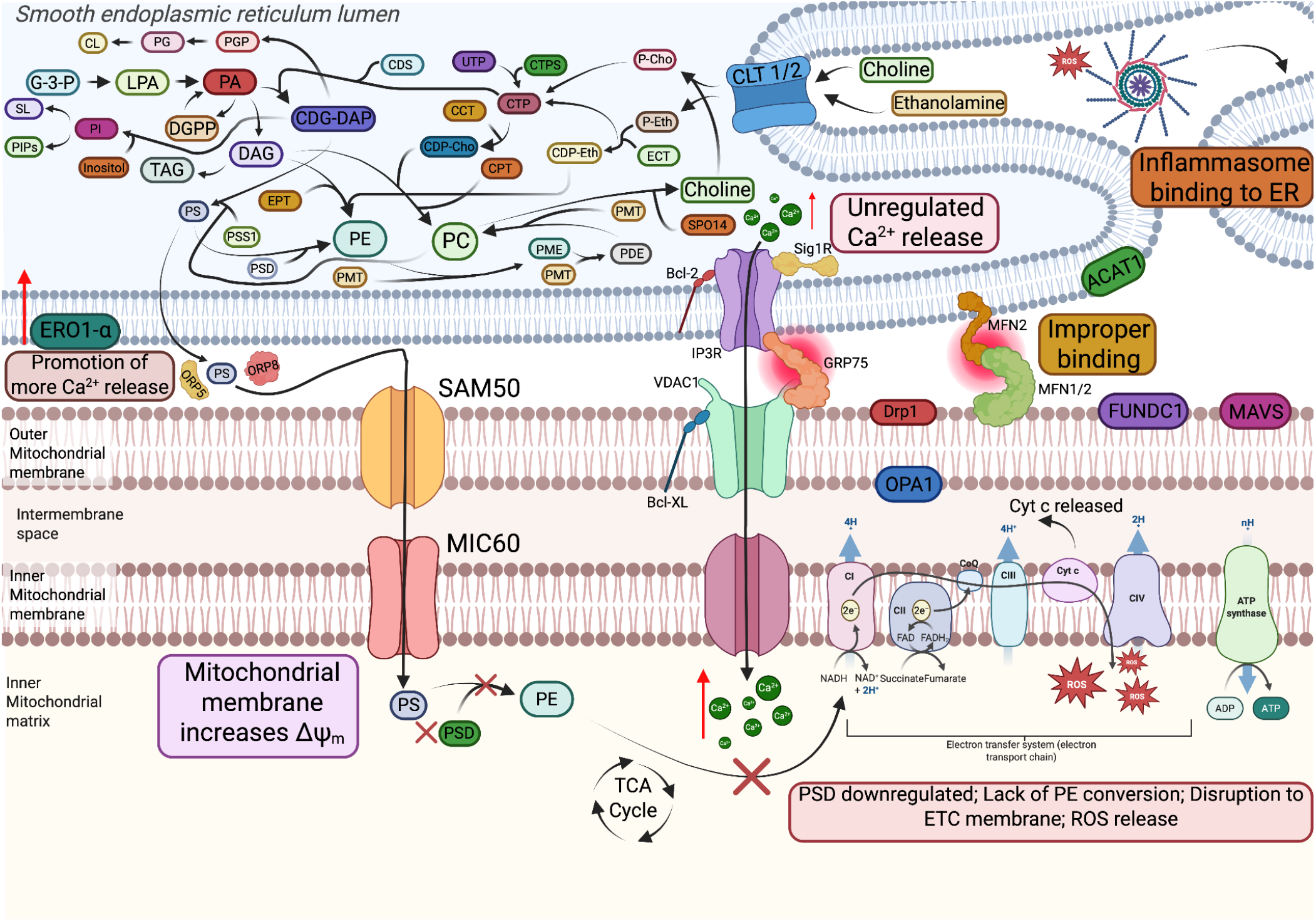
Proposed model of mitochondrial-associated endoplasmic reticulum (ER) membrane (MAM) dysfunction and phospholipid remodeling in spontaneously hypertensive rat (SHR) brain during chronic neuroinflammation. The MAM serves as a critical signaling hub coordinating phospholipid biosynthesis, calcium homeostasis, mitochondrial dynamics, and bioenergetic function through intimate communication with the endoplasmic reticulum (ER) and mitochondria. Chronic hypertensive stress promotes oxidative damage and widespread remodeling of membrane phospholipids, including phosphatidic acid (PA), phosphatidylcholine (PC), phosphatidylenthanolamine (PE), phosphatidylserine (PS), and phosphatidylinositol (PI) disrupting membrane integrity and homeostasis. Upregulation of PS through the SAM50/MIC60 complex combined with the downregulation of phosphatidylserine decarboxylase (PSD) via enzymatic disruption results in a downregulation of PE synthesis from the mitochondrial membrane, a key component in the organelle membrane structure. This destabilizes the inner mitochondrial membrane (IMM) and the electron transport chain, further resulting in sphingolipid reprogramming, loss of phospholipid asymmetry, biosynthetic disruption (exemplified by PDME), and mitochondrial compromise. These interrelated processes converge on vulnerable brain regions, providing a robust mechanistic framework linking hypertension to cognitive decline. By delivering spatially resolved data, integrative modeling (Figure 11), and comprehensive supplementary pathway analysis (Supplementary Figures S1-S11), this study not only deepens our understanding of neuroinflammatory pathology but also identifies actionable therapeutic targets with strong potential to mitigate chronic neuroinflammation-driven brain injury.

The ability of MALDI-MSI to simultaneously map major lipids, oxidized species, biosynthetic intermediates (e.g., PDME), and spatial patterns of vulnerability, and those are complemented by detailed pathway schematics in Supplementary Figures S1-S11. This offers mechanistic resolution far beyond conventional bulk lipidomics. The ODE model (Figure 10) further strengthens this framework by quantitatively reproducing the directional changes and demonstrating bistable-like behavior, whereby sufficiently strong neuroinflammatory input shifts the system into a self-sustaining pathological state.

This study represents a significant advance as one of the first high-resolution spatial lipidomic investigations of hypertensive brain injury. By integrating region-specific multi-omics data with dynamic modeling and mechanistic diagrams, it establishes a detailed lipid-centric model of chronic neuroinflammation that bridges hypertension, mitochondrial dysfunction, and neurodegenerative risk. The findings have broad translational relevance for vascular contributions to cognitive impairment and dementia.

From a therapeutic standpoint, the identified nodes, particularly the oxidative-nSMase2-ceramide-MAM-mitochondrial axis (Figure 11), which offer multiple high-priority targets. Selective nSMase2 inhibitors could break the ceramide-driven vicious cycle; mitochondria-targeted antioxidants or MAM stabilizers could mitigate ROS amplification and Ca^2+^ dysregulation; and strategies supporting phospholipid methylation or membrane asymmetry could preserve cellular resilience. The ODE model enables systematic in silico screening of combination therapies, predicting synergistic efficacy. Chronic neuroinflammation in the SHR model induces a coordinated, MAM-centered membrane remodeling program characterized by advanced oxidative damage, excessive reactive oxygen species (ROS) production. Simultaneously enhancing ERO1-α and dysregulated IP3R-GRP75-VDAC1 signaling promote Ca^2+^ from the ER to the mitochondria. Disrupted MFN1/2-mediated tethering and Drp1 overactivation induce mitochondrial fragmentation, cytochrome c release, inflammasomes binding to the ER, and disrupted oxidative phosphorylation further upregulate the inflammatory response resulting in an upregulation of ceramide, and later, sulfatide lipid remodeling.Together, these alterations establish an upregulation of mitochondrial dysfunction and oxidative stress that amplifies phospholipid remodeling, inflammatory signaling, and membrane damage, providing a mechanistic framework linking MAM dysfunction to chronic neuroinflammation, white matter injury, and progressive neurodegeneration in the SHR brain.

## Conclusion

In conclusion, this spatial lipidomic study using positive-ion mode MALDI-MSI demonstrates that chronic neuroinflammation in the SHR hypertensive brain triggers a profound, coordinated multi-class membrane remodeling program dominated by advanced lipid peroxidation, nSMase2-driven sphingolipid dysregulation, loss of phospholipid asymmetry, and mitochondrial bioenergetic compromise. These interconnected processes, mechanistically integrated through the nSMase2–ceramide–Drp1–ROS–NF-κB vicious cycle (Figure 11) and quantitatively supported by ODE modeling (Figure 10), converge on vulnerable regions such as the hippocampus and white-matter tracts, providing a lipid-based mechanistic bridge between hypertension, neuroinflammation, and heightened risk of cognitive decline and Alzheimer’s disease pathology. By combining high-resolution spatial mapping with pathway-focused mechanistic schematics and dynamic modeling, the work identifies the oxidative–sphingolipid–mitochondrial axis as a high-value therapeutic target. Targeting this axis—particularly through nSMase2 inhibition, mitochondria-directed antioxidants, and membrane stabilization strategies—holds strong promise for interrupting chronic neuroinflammatory cascades, not only in hypertensive brain injury but also across a spectrum of age-related neurodegenerative conditions. This study thus advances both mechanistic understanding and translational opportunities for mitigating neuroinflammation-driven brain pathology.

### Limitations

Several limitations should be considered. First, the analysis was performed exclusively in positive-ion mode with DAN matrix, which provides excellent coverage of PCs, SMs, HexCers, and ceramides but offers limited detection of anionic phospholipids (e.g., certain PS, PI, and PA species in negative-ion mode). Complementary negative-ion mode imaging would provide a more complete lipidomic picture. Second, while the SHR model effectively recapitulates chronic hypertension and associated neuroinflammation, findings require validation in additional models and, ultimately, in human hypertensive and VCI/AD brain tissues. Third, the study focuses on end-stage lipidomic signatures; longitudinal analyses at earlier time points would further clarify the temporal evolution of these changes and the point at which interventions could most effectively halt progression. Finally, although the ODE model successfully reproduces the observed directional changes and bistable behavior, it represents a simplified abstraction of the underlying biology and will benefit from iterative refinement with additional quantitative parameters (e.g., regional antioxidant capacity or specific enzyme kinetics). Despite these limitations, the spatially resolved insights and mechanistic framework presented here provide a solid foundation for future hypothesis-driven research and therapeutic development targeting chronic neuroinflammation.

## Notes

The authors declare no competing financial interests.

## ACKNOWLEDGMENT

We sincerely acknowledge financial support from the NIH (R01HL163159, Z.S.; R15 EB035866, L.B.), the American Heart

Association (AHA grant #1807047, L.B.), and GLRC-ICC (R01805, L.B.). All authors, especially the graduate students, gratefully acknowledge the Department, Dr. Will Cantrell, and the Graduate School for their support of student training and research progress. We also extend our deep gratitude to Dr. Rick Koubek for his encouragement and unwavering support throughout this project.

