## Supplemental Material for "Spatial MALDI-MSI Reveals a Coordinated Vicious Cycle of Oxidative Membrane Damage and Ceramide-Driven Sphingolipid Dysregulation in the Chronically Neuroinflamed Brain"

**ASSOCIATED CONTENT**

**Supporting Information**

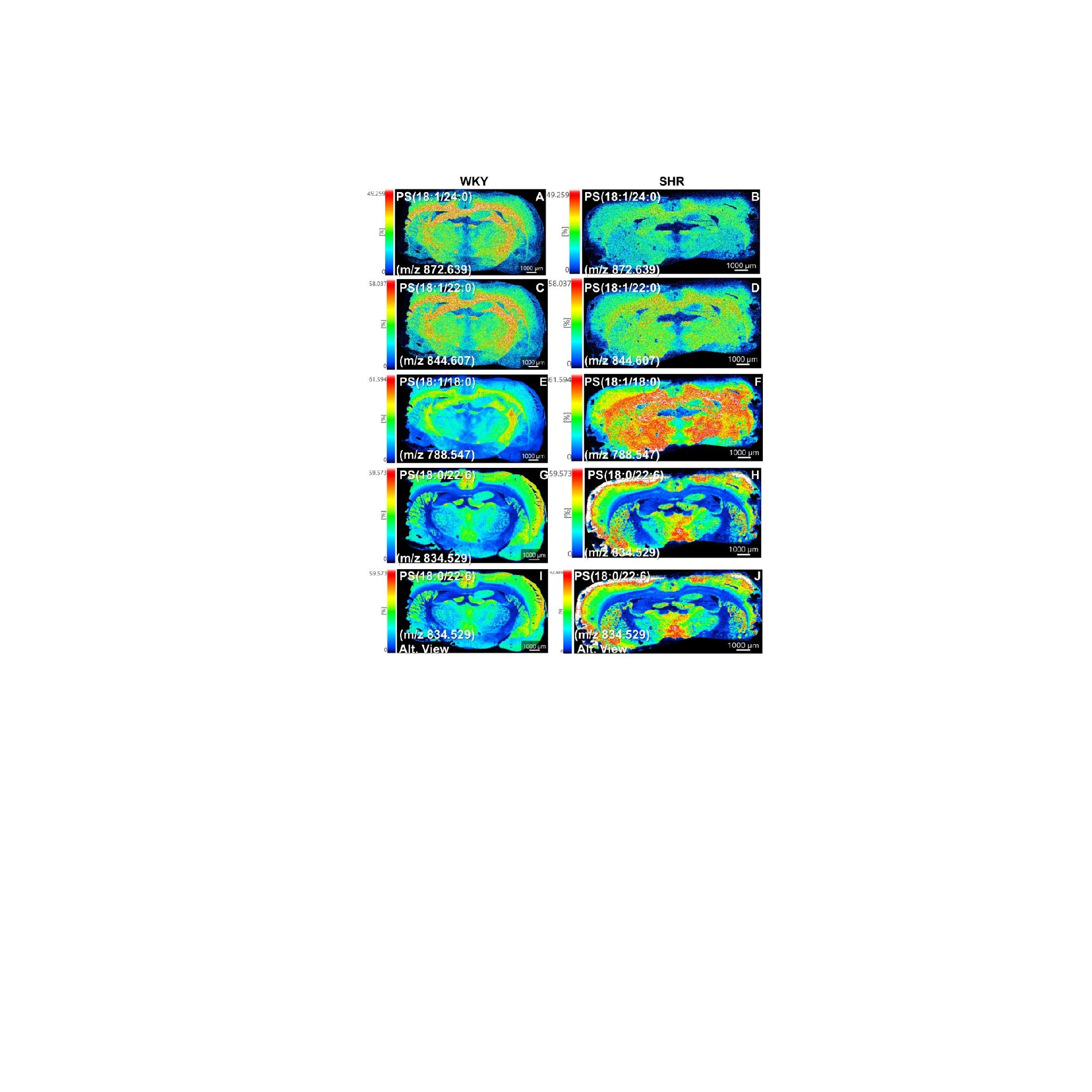

Figure S1. Additional phosphatidylserine (PS) species in WKY normotensive (left column) and SHR hypertensive (right column) rat brain sections. Ion images of long-chain and polyunsaturated PS species not shown in main Figure 4.. Color scale represents relative ion intensity (% of maximum, normalized across all panels). Scale bar = 1000 μm. This supplementary figure expands the documentation of PS distribution patterns and supports the finding of increased regional heterogeneity and membrane remodeling in the SHR hypertensive brain (see main Figure 4)

**S1.1. Additional Phosphatidylserine Species and Their Spatial Distribution in WKY and SHR Brains**

Supplementary Figure S1 presents additional and alternative phosphatidylserine (PS) panels.. The selected species include long-chain and polyunsaturated PS, specifically PS(18:1/24:0) at *m/z* 872.6386, PS(18:1/22:0) at *m/z* 844.6073, PS(18:1/18:0) at *m/z* 788.5470, PS(18:0/22:6) at *m/z* 834.5291, and PS(18:1/22:6) at ~*m/z* 834.5.

In WKY normotensive control brains, these PS species generally displayed relatively uniform and homogeneous distribution across cortical and hippocampal regions. In contrast, SHR hypertensive brains exhibited more heterogeneous and patchy signal patterns, with noticeable variations in intensity between different cortical layers and hippocampal subregions. These spatial differences were reproducible across multiple PS species and views.

**S1.2. Enhanced Regional Heterogeneity of PS Signals in the SHR Hippocampus**

A particularly notable feature in SHR sections was the increased regional heterogeneity and focal signal patterns of PS species within the hippocampus. While WKY controls showed relatively even PS distribution, SHR brains frequently displayed discrete areas of higher or lower intensity, creating a patchy appearance. This heterogeneity was especially evident in long-chain species such as PS(18:1/24:0) and polyunsaturated species such as PS(18:0/22:6). The use of both paired images and a full single SHR panel (Row 5) further confirmed the robustness of these regional differences.

**S1.3. PS Heterogeneity as Evidence of Membrane Remodeling and Apoptotic Signaling**

The additional PS panels in Supplementary Figure S1 provide further support for hypertension-associated membrane remodeling. Phosphatidylserine is normally maintained on the inner leaflet of the plasma membrane. Its redistribution or altered spatial organization is an early indicator of loss of membrane asymmetry and is closely linked to apoptotic signaling, as externalized PS serves as a recognition signal for microglial phagocytosis of damaged cells.

The more heterogeneous and patchy PS distribution observed in SHR brains, particularly in the hippocampus, is consistent with increased localized membrane stress and apoptotic activity under chronic hypertensive conditions. These findings align with the oxidative stress and lipid peroxidation documented in Figures 1 and 2, which can disrupt membrane integrity and promote PS redistribution. The hippocampal accentuation is biologically significant given the high vulnerability of this region to hypertensive injury and its central role in cognitive function.

**S1.4. Integration with Main Figure 4 and Overall Mechanistic Implications**

When combined with the PS findings in main Figure 4, Supplementary Figure S1 strengthens the conclusion that hypertension induces regionally heterogeneous membrane phospholipid remodeling, with the hippocampus showing particular vulnerability. The additional views of long-chain and polyunsaturated PS species confirm that the increased heterogeneity is not limited to a few species but is a consistent feature across the PS class.

These PS alterations complement the oxidative damage to phosphatidylcholines (Figures 1–2), sphingolipid and ceramide dysregulation (Figures 3 and 9), and changes in other phospholipid classes (Figures 5–8). Together, they support a model in which chronic hypertensive stress drives coordinated membrane injury involving lipid peroxidation, loss of asymmetry, and apoptotic signaling. The spatially resolved PS heterogeneity provides direct evidence that membrane remodeling in the hypertensive brain is region-specific and likely contributes to neurovascular dysfunction and long-term cognitive impairment.

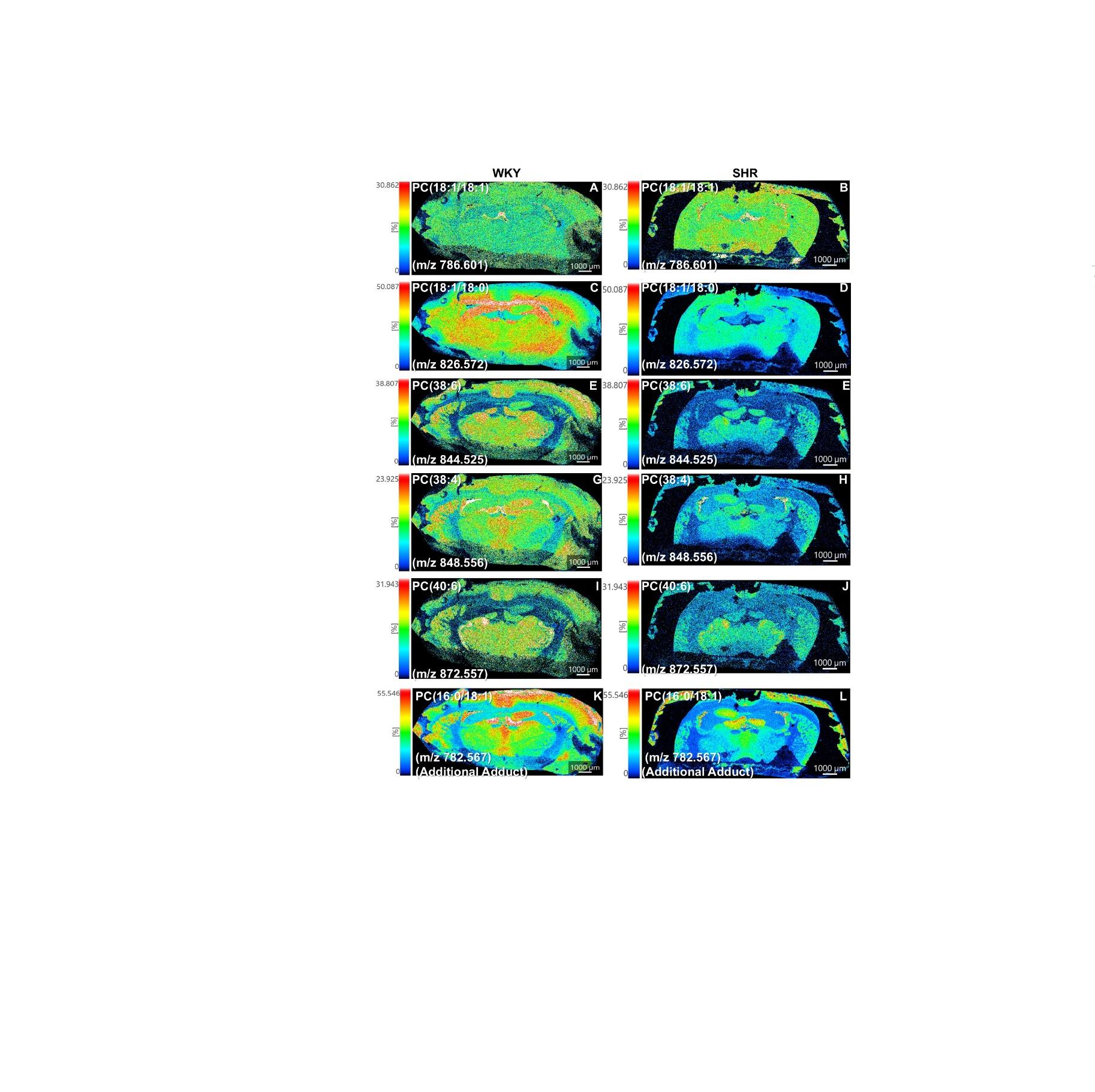

Figure S2. Additional phosphatidylcholines (PC) in WKY normotensive and SHR hypertensive rat brain sections (complete class coverage). This supplementary figure presents the remaining high-quality PC panels that were not included in main Figure 2, providing full documentation of the phosphatidylcholine class to support the oxidative stress and membrane remodeling findings.

S2.1. Additional Phosphatidylcholine Species and Their Spatial Distribution

Supplementary Figure S2 presents additional phosphatidylcholine (PC) panels. In WKY normotensive control brains, these PC species generally exhibited relatively strong and widespread signals across both gray and white matter regions. In contrast, SHR hypertensive brains showed consistently reduced signal intensity for most of these structural PC species. The spatial distribution in SHR sections appeared more attenuated, particularly in white-matter tracts and subcortical areas, compared with the robust and diffuse distribution observed in WKY controls.

S2.2. Quantitative Trends Supporting Class-Wide Downregulation

Region-of-interest analysis and visual comparison across the additional PC panels confirmed a consistent trend of reduced abundance in SHR brains. Long-chain polyunsaturated species such as PC(38:6), PC(38:4), and PC(40:6) showed moderate to marked reductions in signal intensity. Similarly, PC(18:1/18:0) [M+K]⁺ and PC(18:0/22:6) exhibited lower signals in SHR sections. The additional adduct panels (Na⁺ and K⁺ forms of PC(16:0/18:1) and PC(16:0/16:0)) further validated that the downregulation of major diacyl PCs is reproducible across different ionization states.

**S2.3. Data Interpretation: Support for Broad Membrane Phospholipid Remodeling**

The additional PC species presented in Supplementary Figure S2 strengthen the conclusion that hypertension induces a broad depletion of major structural phosphatidylcholines. The consistent reduction observed across diacyl, long-chain, and polyunsaturated PC species indicates that membrane phospholipid remodeling in the SHR brain is not limited to a few specific molecules but affects the PC class in a widespread manner.

This class-wide downregulation of structural PCs complements the pronounced upregulation of oxidized and short-chain PCs documented in main Figure 2. Together, these reciprocal changes suggest that chronic hypertension promotes both the generation of oxidized lipid species through lipid peroxidation and the loss or accelerated turnover of abundant membrane phospholipids. Such coordinated remodeling likely compromises membrane integrity, fluidity, and the organization of membrane microdomains, thereby affecting cellular signaling and mitochondrial function.

**S2.4. Integration with Main Figure 2 and Overall Lipidomic Findings**

When combined with the findings from main Figure 2, Supplementary Figure S2 provides complete class coverage of phosphatidylcholines and confirms that the pattern of oxidative PC accumulation accompanied by structural PC depletion is a robust and reproducible feature of the SHR hypertensive brain. The additional panels demonstrate that this remodeling extends across multiple molecular species and adduct forms, reinforcing the biological significance of these changes.

These PC alterations integrate with the sphingolipid dysregulation (Figure 3 and Figure 9), the heterogeneous distribution of PS, PI, and PE species (Figures 4–6), and the changes in minor phospholipids and metabolites (Figures 7 and 10). Collectively, they support a model in which hypertension drives multi-class membrane phospholipid remodeling characterized by oxidative damage to structural lipids and disruption of membrane homeostasis. The spatially resolved nature of these changes, particularly the attenuation of structural PCs in white-matter and subcortical regions, provides further evidence that membrane injury contributes to the neurovascular and cognitive consequences of chronic hypertension.

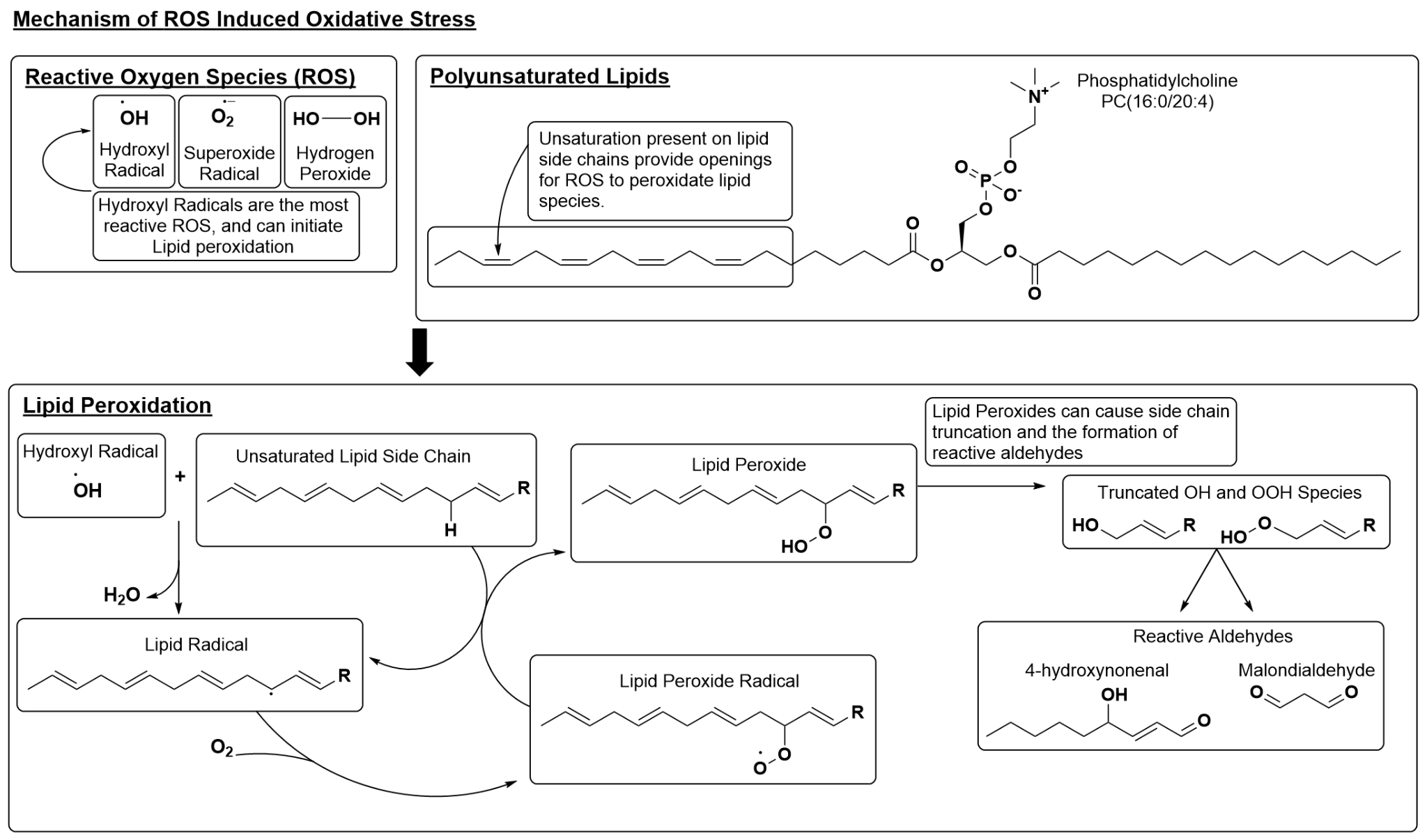

Figure S3. Mechanism of ROS-induced lipid peroxidation. Reactive oxygen species (ROS), particularly hydroxyl radicals, abstract bis-allylic hydrogens from polyunsaturated fatty acyl chains in membrane phospholipids. This initiates lipid peroxidation, generating lipid radicals that propagate to form lipid hydroperoxides and, ultimately, truncated oxidized phospholipids and reactive aldehydes (e.g., 4-hydroxynonenal). The scheme illustrates the key chemical steps and downstream consequences relevant to oxidative membrane damage in the SHR brain.

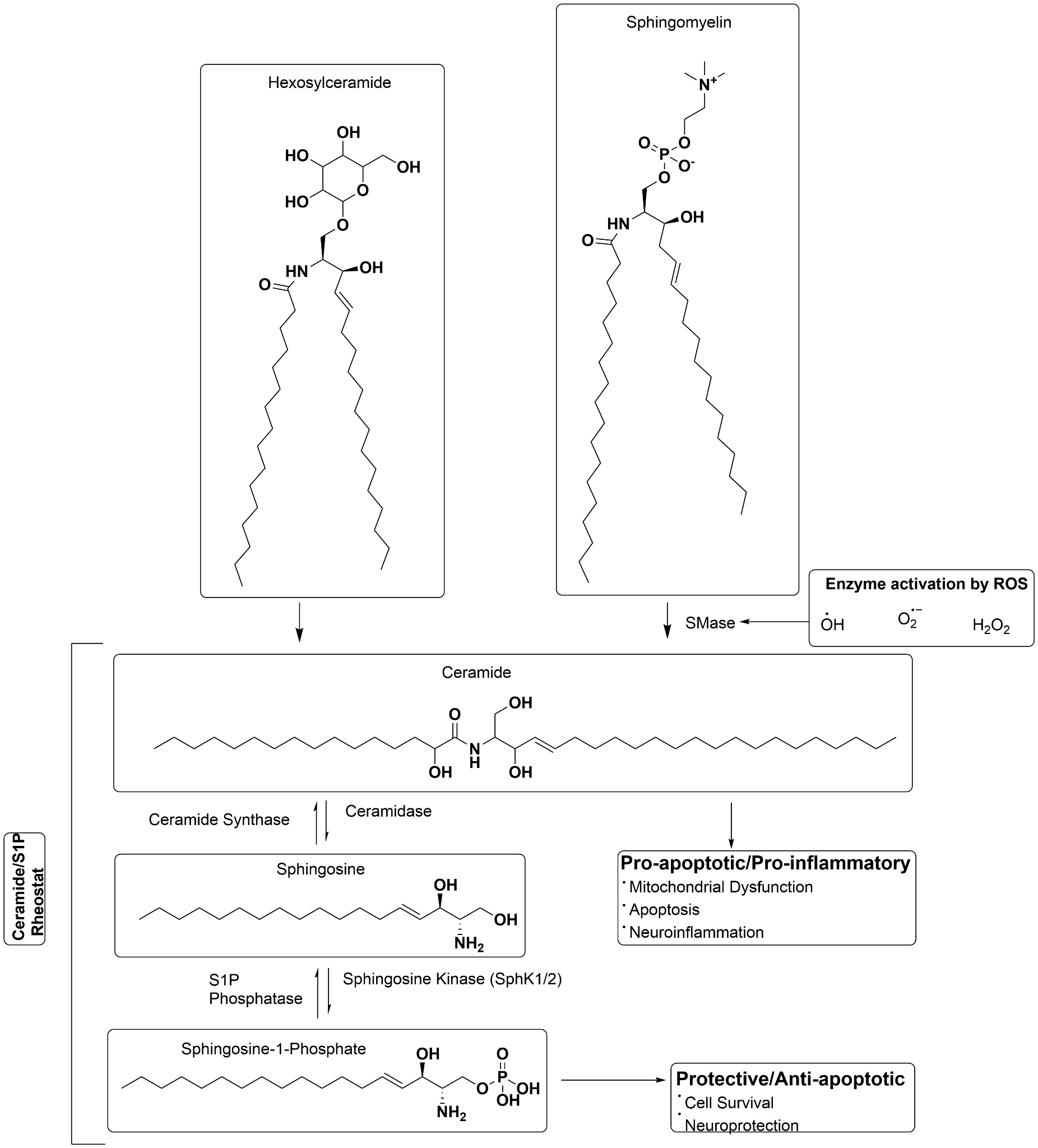

Figure S4. Sphingolipid metabolism and the ceramide/S1P rheostat in the SHR brain. Upregulation of sphingomyelin and hexosylceramide species, driven by ROS-activated neutral sphingomyelinase (nSMase2), promotes ceramide accumulation. The resulting shift in the ceramide/S1P rheostat favors pro-apoptotic and pro-inflammatory signaling (mitochondrial dysfunction, apoptosis, endothelial impairment, and neuroinflammation) over protective S1P-mediated pathways.

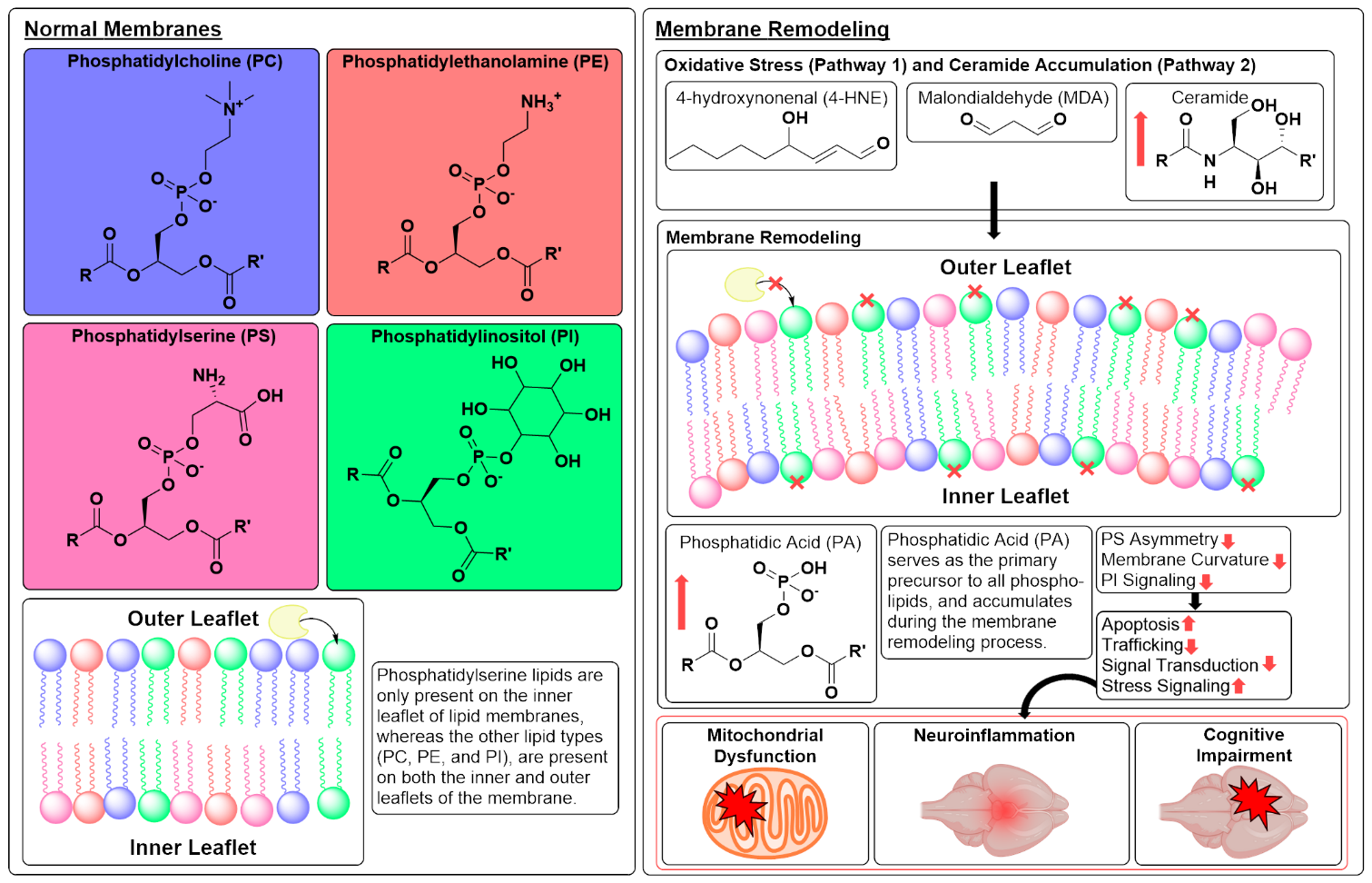

Figure S5. Phospholipid asymmetry and membrane remodeling during oxidative stress. In healthy membranes, phosphatidylserine (PS) is restricted to the inner leaflet while phosphatidylinositol (PI) species serve as signaling molecules whose function depends on phosphorylation of the inositol headgroup. Oxidative stress and ceramide accumulation disrupt asymmetry, alter membrane curvature, and impair signaling, promoting apoptosis and neuroinflammation (right panel).

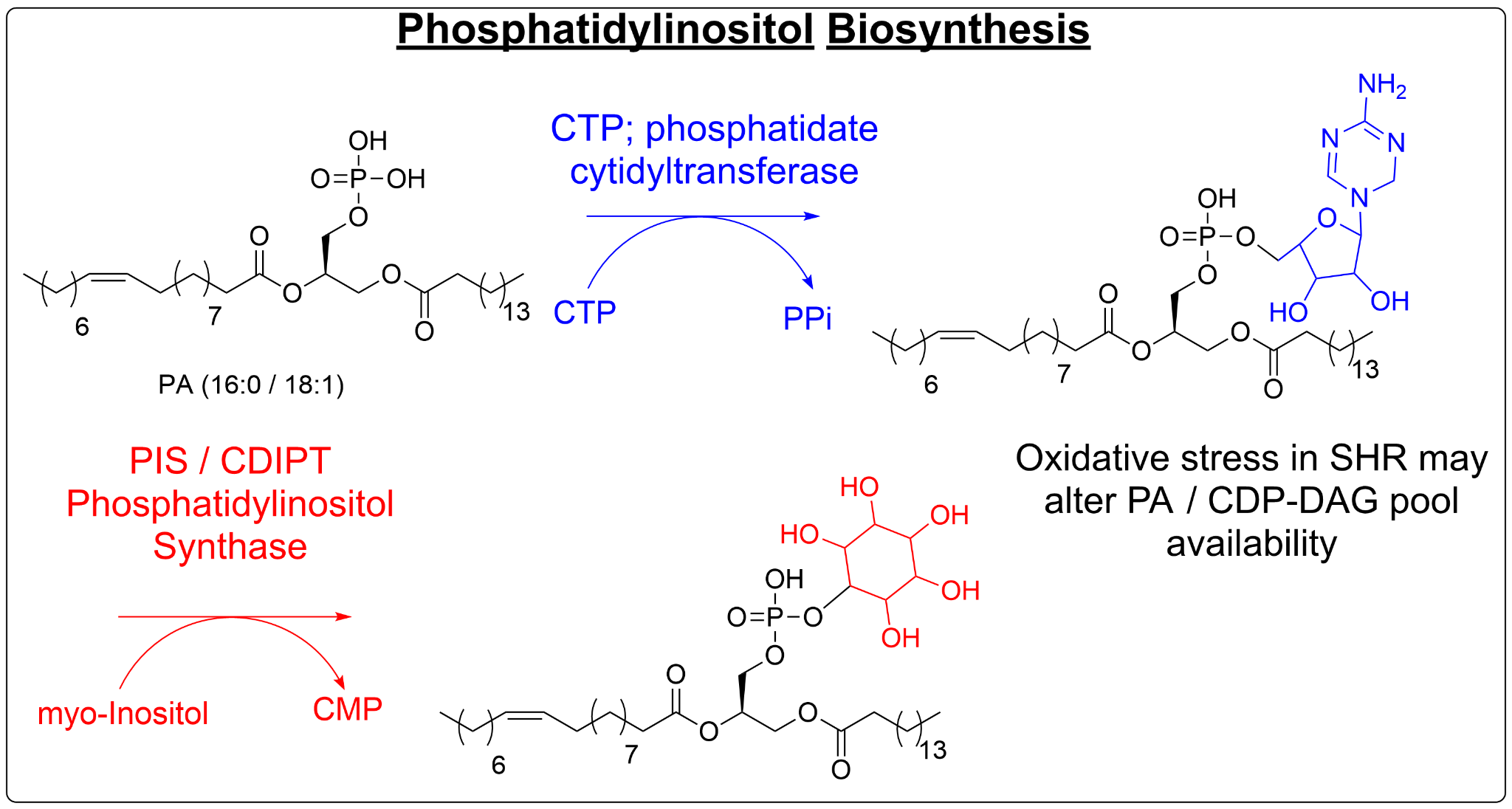

Figure S6. Phosphatidylinositol biosynthesis pathway. Phosphatidic acid (PA) is converted to CDP-diacylglycerol by phosphatidate cytidylyltransferase (CTP:PA cytidylyltransferase). Phosphatidylinositol synthase then combines CDP-DAG with myo-inositol to form phosphatidylinositol (PI). Oxidative stress in the SHR brain may alter PA availability and thereby influence PI signaling and membrane properties.

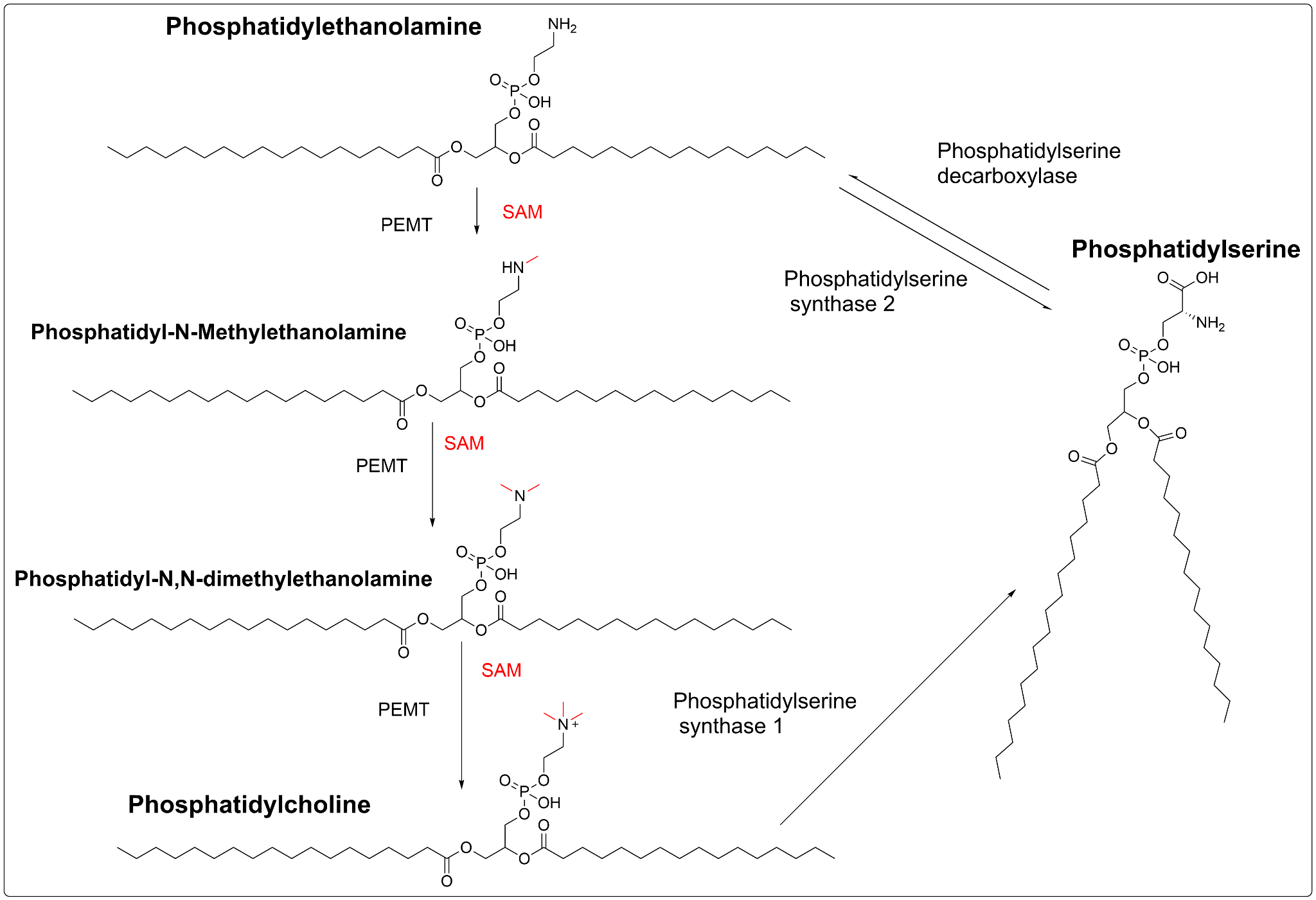

Figure S7. Biosynthesis of phosphatidylcholine via the methylation pathway and conversion to phosphatidylserine. Phosphatidylethanolamine (PE) is sequentially methylated by phosphatidylethanolamine N-methyltransferase (PEMT) using S-adenosylmethionine (SAM) to form phosphatidyl-N-methylethanolamine, phosphatidyl-N,N-dimethylethanolamine (PDME), and ultimately phosphatidylcholine (PC). Both PC and PE can be converted to phosphatidylserine (PS) by phosphatidylserine synthases 1 and 2. Accumulation of PDME in the SHR brain suggests dysregulation of this pathway under chronic neuroinflammatory stress.

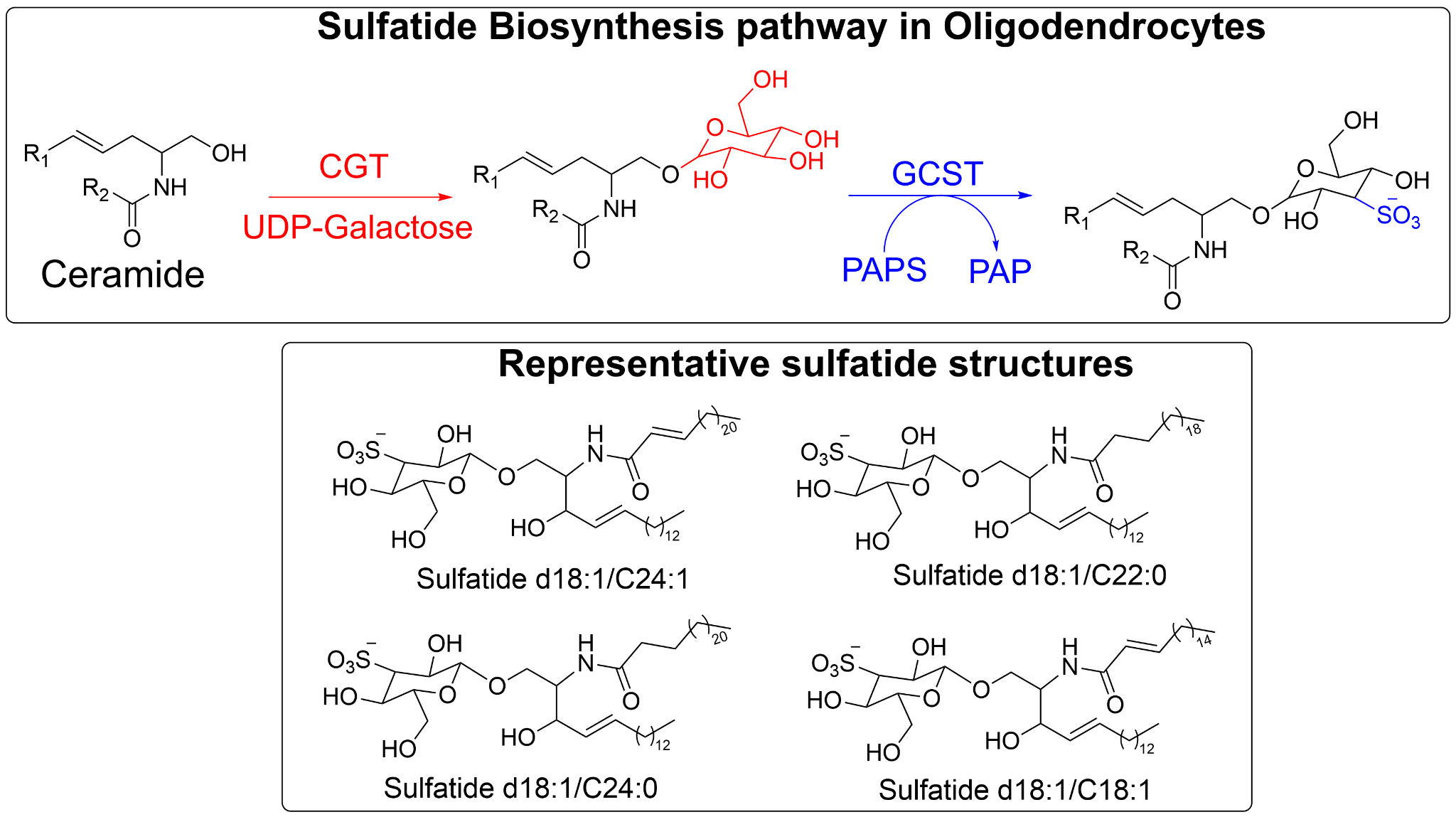

Figure S8. Sulfatide biosynthesis pathway in oligodendrocytes. Ceramide is converted to galactosylceramide by ceramide galactosyltransferase (CGT) using UDP-galactose, then sulfated by cerebroside sulfotransferase (CST/GCST) with 3'-phosphoadenosine-5'-phosphosulfate (PAPS) to form sulfatide. Representative sulfatide structures with common acyl chain lengths are shown. Alterations in this pathway contribute to myelin remodeling observed in the SHR white matter.

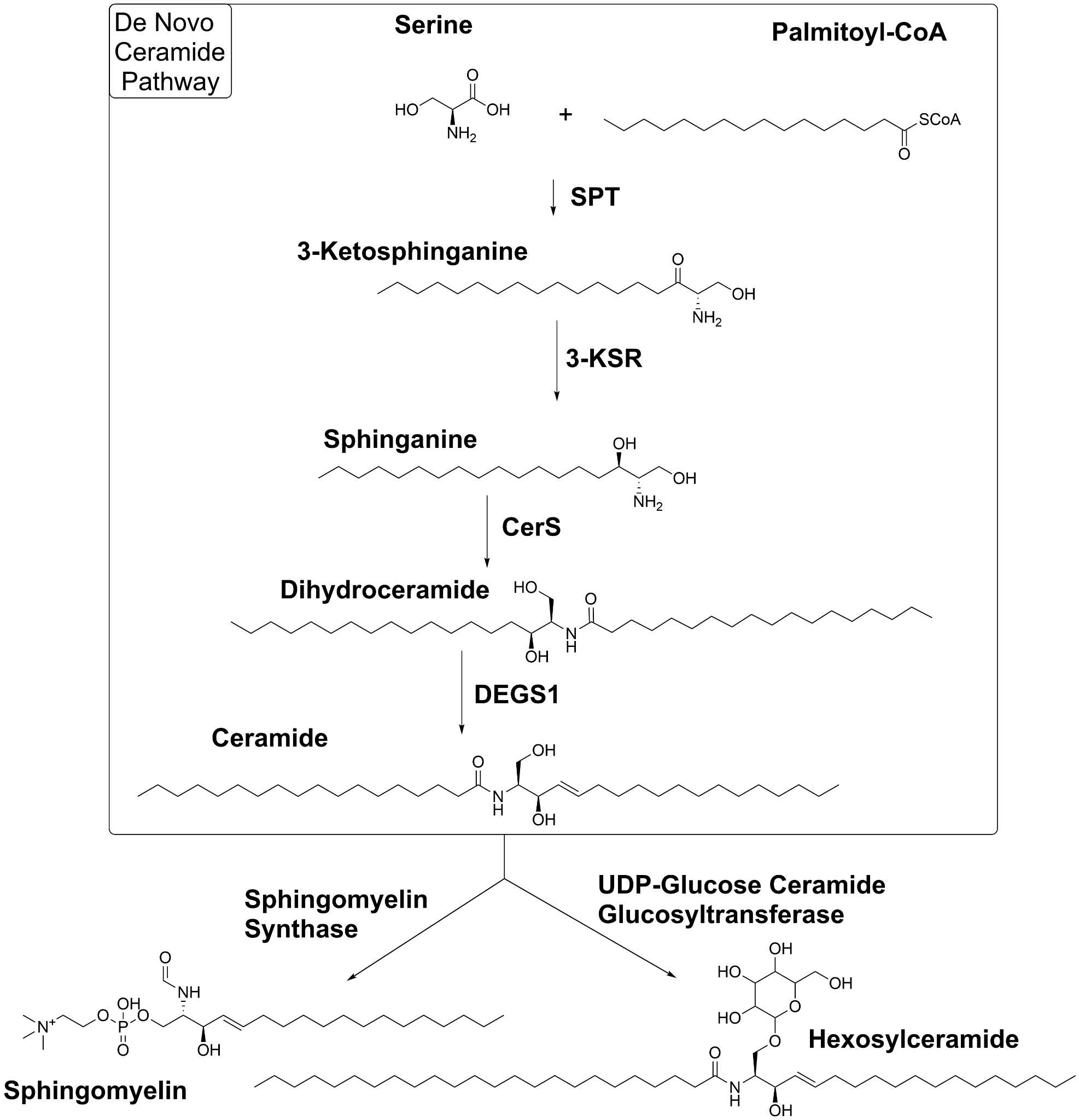

Figure S9. De novo ceramide synthesis and conversion to sphingomyelin and hexosylceramide. Serine and palmitoyl-CoA are condensed by serine palmitoyltransferase (SPT) to form 3-ketosphinganine, which is reduced to sphinganine. Acylation by ceramide synthase (CerS) followed by desaturation (DEGS1) yields ceramide. Ceramide is then converted to sphingomyelin via sphingomyelin synthase or to hexosylceramide via UDP-glucose ceramide glucosyltransferase. This pathway supports the coordinated sphingolipid upregulation observed in the SHR brain.

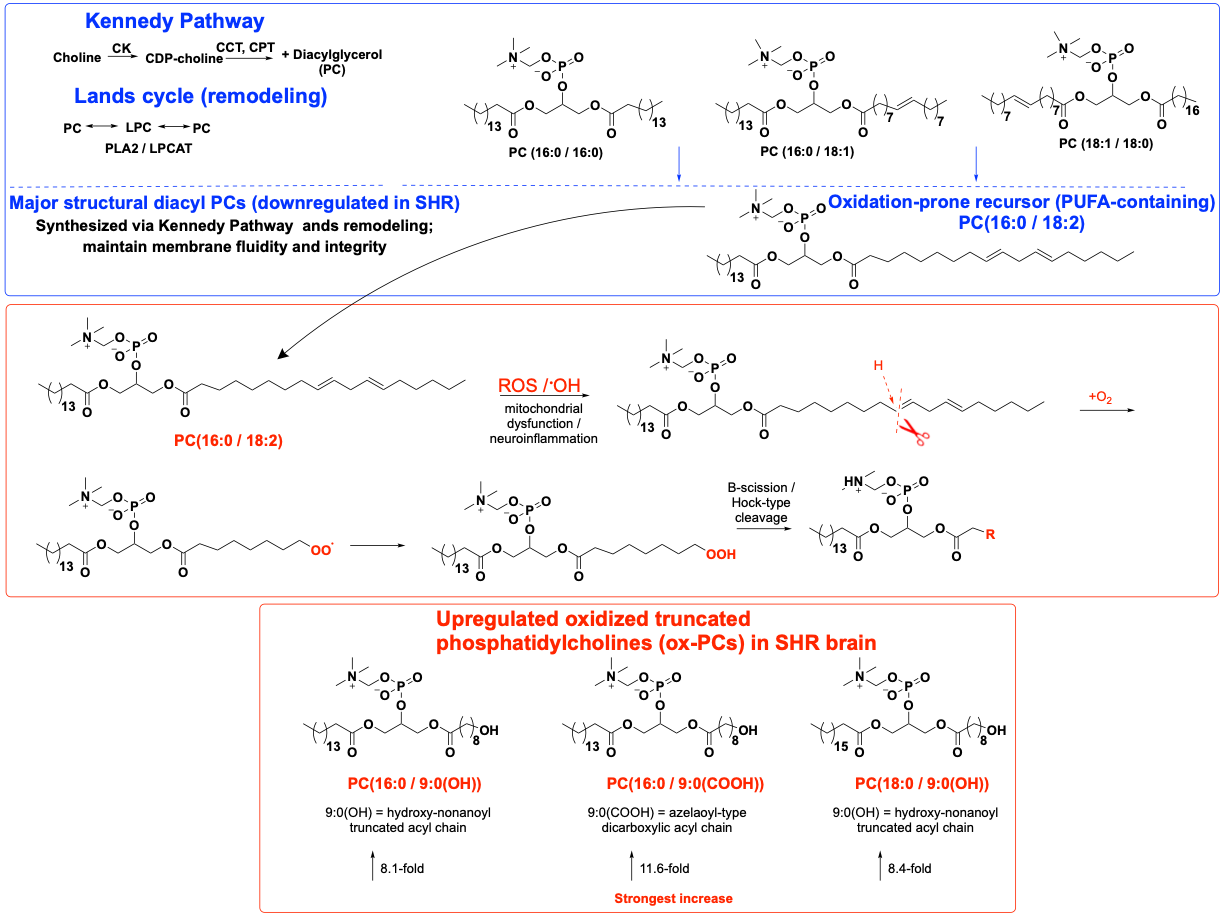

Figure S10. Integration of phospholipid biosynthesis, remodeling, and oxidative truncation in the SHR brain. The Kennedy and Lands cycles maintain major structural diacyl PCs (downregulated in SHR). ROS attack on PUFA-containing precursors drives lipid peroxidation and chain truncation, generating upregulated oxidized PCs (e.g., PC(16:0/9:0(COOH)), PC(16:0/9:0(OH)), PC(18:0/9:0(OH))). The schematic summarizes how chronic neuroinflammation simultaneously impairs structural phospholipid homeostasis and promotes formation of bioactive truncated oxidized species.

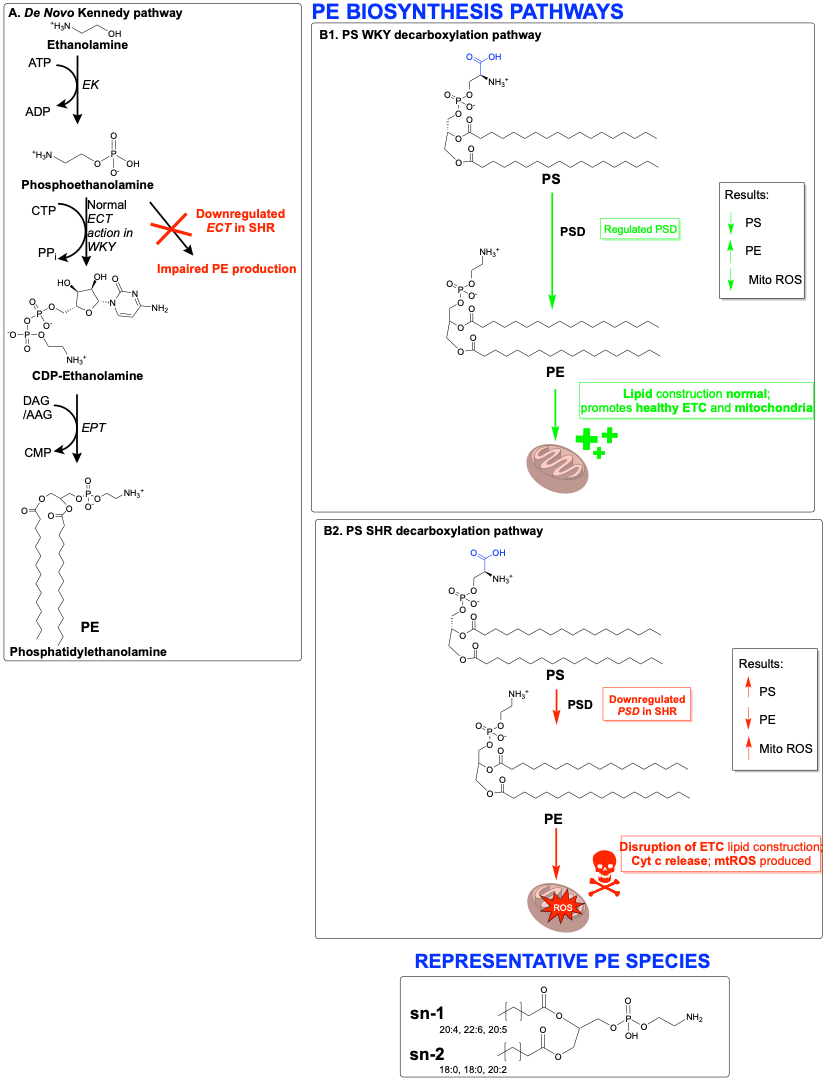

Figure S11. Biosynthesis of PE in both WKY and SHR. The regulation of PSD in WKY rats results in an upregulation of PE which makes up the majority of the electron transport chain's lipid construction. An upregulation of PE results in an upregulation of healthy mitochondria. In SHR rats, PSD is down regulated. This results in a downregulation of PE being converted from PS. PS buildup in the mitochondria results in disruptions to the electron transport chain, consequently releasing cytochrome C. Once cytochrome c is released, this results in an overproduction of mtROS, leading to caspase -9 and -3, further leading to apoptosis.
